# The history of enzyme evolution embedded in metabolism

**DOI:** 10.1101/2025.07.16.665256

**Authors:** Tatsuya Corlett, Harrison B. Smith, Eric Smith, Joshua E. Goldford, Liam M. Longo

## Abstract

Whereas phylogenetic reconstructions are a primary record of protein evolution, it is unknown whether the deep history of enzymes are encoded at higher levels of biological organization. Here, we demonstrate that the emergence and reuse history of enzymatic folds is embedded within the web of metabolite-cofactor-enzyme interdependencies that comprise biosphere-scale metabolic reaction networks. Using a simple network analysis approach, we reconstruct the relative ordering of enzymatic fold emergence and, where possible, the first reaction(s) that each enzymatic fold catalyzed. We find that a large majority of enzymatic folds were sufficient as independent additions to open new avenues for metabolic growth. The resulting network-based histories are broadly concordant with enzyme phyletic distribution in prokaryotes, a proxy for enzyme age. Our results suggest that the earliest enzyme-mediated metabolisms were enriched for α/β proteins, likely due to their strong association with cofactor utilization, and that α-proteins preferentially emerge at later stages. The cradle-loop barrel, a member of the small β-barrel metafold, is predicted to be the founding β-fold, in agreement with analyses of ribosome structure. An examination of how the protein universe responded to the biological production of molecular oxygen reveals that the adaptation of existing enzymatic folds, not novel fold emergence, was the primary driver of metabolic evolution. This work presents a self-consistent model of metabolic and enzyme evolution, key progress towards integrating diverse perspectives into a unified history of protein evolution.

**Significance Statement:** Enzyme emergence is an ongoing process that began ∼4 billion years ago. Here, we show that the modern biosphere-scale network of metabolic reactions and enzymes is an archive of enzyme history independent from, but concordant with, phylogenetics. Based on this record, we predict the order of enzyme emergence from before the last universal common ancestor up until the biological production and metabolic utilization of molecular oxygen. We find that while α/β proteins dominated primitive enzyme-mediated metabolism, other folds — including the cradle-loop barrel, which is a member of the small β-barrel metafold — were likely important early contributors. This study represents key progress towards building an internally consistent, joint history of metabolic reactions and enzymes.

## Introduction

The last universal common ancestor (LUCA) possessed a repertoire of complex protein domains <u>(Moody et al. 2024)</u>, indicating that key events in protein evolution occurred even earlier in the history of life. While extant protein sequences are a rich archive of protein evolution, understanding protein fold emergence and evolution prior to the LUCA poses a unique challenge <u>(Winstanley et al. 2005; Edwards et al. 2013)</u>. Because the LUCA represents the approximate point at which historical inference by comparative approaches converges, resolution at the earliest stages of protein evolution is poor <u>(Winstanley et al. 2005; Edwards et al. 2013)</u>. Thus, alternative records of protein evolution are needed to reconstruct the fold emergence process. What records may be suitable for this purpose?

One alternative record of early protein evolution that has been proposed is the ribosome, in which domains or pieces of domains have been “frozen in time” by accretionary processes <u>(Petrov et al. 2015; Kovacs et al. 2017)</u>. By analyzing the layered structure of the ribosome, an early history of protein evolution can be inferred. This approach indicates the early emergence of β-structural elements prior to other structure classes — a surprising result, given that β-protein composition within proteomes increases markedly after the emergence of chaperones <u>(Rebeaud</u> <u>et al. 2021)</u>, perhaps suggesting that β-proteins have intrinsic deficits to foldability <u>(Plaxco et al.</u> <u>1998; Takács et al. 2024)</u>. Although the extent to which ribosomal proteins inform the emergence of other protein folds is unknown, this work represents a remarkable insight into the types of pre-LUCA archives that may record the earliest stages of protein evolution.

Another layered system in biology is the structure of metabolism <u>(Handorf et al. 2005;</u> <u>(Raymond and Segrè 2006)</u>. Whereas physical stratification suggests an emergence order in the ribosome, metabolic layering results from the network of substrate, cofactor, and enzyme requirements for each metabolic reaction. Viewing metabolism through this lens has produced significant insights into the chemistry of primitive metabolic systems <u>(Goldford et al. 2017)</u>, the environment of the earliest life <u>(Goldford et al. 2019)</u>, and the existence of forgotten chemical reactions <u>(Goldford et al. 2024)</u>. We note that these prior studies were independent of enzyme history, leaving a gap in understanding about the co-evolutionary history of metabolic networks and enzymes. We hypothesize that the early history of enzyme emergence and evolution may also be written into the web of known metabolic interdependencies.

Here, we reconstruct the order of enzyme evolution based on the layered structure of metabolism using a novel network analysis algorithm that we refer to as *enzyme-gated network expansion*. We demonstrate that the complex, multifaceted associations between metabolites, reactions, cofactors, and enzymes can constrain the order of enzyme discovery. We find that the handoff from geochemistry-mediated to enzyme-mediated metabolic exploration soon after the availability of nucleotides yields a robust enzyme emergence trajectory with a consistent and realistic progression in enzyme complexity. Our reconstructed order of enzymatic fold emergence is broadly concordant with phylogenetic information. The results suggest an intriguing transition from easily discovered α/β domains to those dependent on more refined evolutionary information and control systems. Using the biological production of molecular oxygen as an example of metabolic upheaval, we characterize the interplay between enzyme reuse — in which a previously discovered enzyme performs a new function — and the emergence of novel enzymatic folds. This work represents a key advancement toward developing a fully integrated history of protein evolution within the framework of a self-consistent model of metabolic evolution.

## Materials and Methods

### Connecting protein evolution and metabolism

Many, though not all, metabolic reactions are catalyzed by enzymes. Enzymes are composed of structural units called protein domains (often several) that can be shared across multiple enzyme families, resulting in complex patterns of partial homology between enzymes. The definition of a protein domain is not fully agreed upon, but is generally meant to entail structural separation, reuse, and independent foldability <u>(Richardson 1981)</u>. Consequently, protein domains are a fundamental unit of protein evolution <u>(Lo Conte et al. 2000; Knudsen and Wiuf 2010; Cheng et al. 2014)</u>. To reconstruct the history of enzyme evolution from metabolism, we must first connect each metabolic enzyme with both the reactions that it catalyzes and the protein domains that comprise its structure (**Figure 1A**).

**Figure 1.**
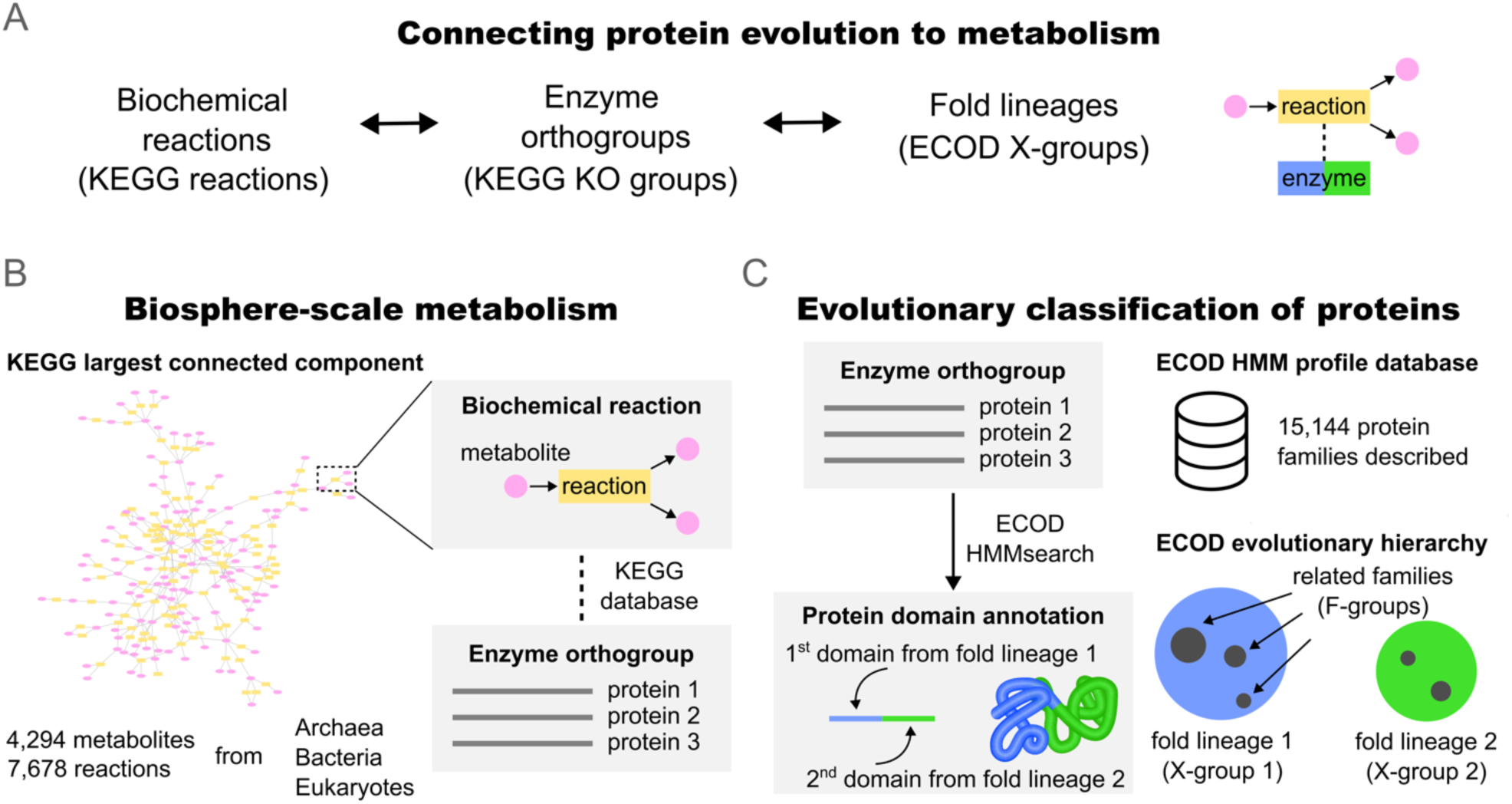
Connecting protein evolution to metabolism. **A**. To simulate the coevolution of metabolism and proteins, biochemical reactions (KEGG reactions) and fold lineages (ECOD X-groups) were connected through mutual association with enzyme orthogroups (KEGG KO groups). **B**. Our metabolic model contains metabolites and reactions encompassing organisms from all branches of the tree of life. Reactions within this network were linked to enzyme orthogroups (KO group) using the manually curated associations within the KEGG database. **C**. For each enzyme orthogroup in KEGG, the evolutionary classifications of the protein domains were assigned based on the ECOD hierarchy. Protein domains can be classified into protein families (F-groups) that are associated with an HMM profile. Protein families can be grouped into fold lineages (X-groups; similar to “clades” in Pfam) that are hypothesized to have emerged from a common founder protein. See **Methods** for detailed explanation of each step.

Connecting enzymes to the reactions that they catalyze has been a longstanding challenge in molecular biology <u>(Sorokina et al. 2014; Hadadi et al. 2019)</u>. The Kyoto Encyclopedia of Genes and Genomes (KEGG) database <u>(Kanehisa and Goto 2000)</u> is a manually curated database of experimentally validated enzyme-reaction associations (**Figure 1B**). The KEGG database includes reactions and enzymes from archaea, bacteria, and eukaryotes, and diverse metabolic strategies are represented <u>(Kanehisa and Goto 2000)</u>. For example, the KEGG database includes the reactions for the seven major carbon fixation pathways. For these reasons, KEGG is considered a repository of biosphere-scale metabolism <u>(Raymond and Segrè 2006)</u>. Here, we employ a previously described model of biosphere-level metabolism based on the KEGG database <u>(Goldford et al. 2024)</u>. In total, our metabolic model <u>(Goldford et al. 2024)</u> comprises 4,294 metabolites and 7,678 reactions. These reactions are associated with 4,331 enzyme orthologous groups, referred to as KO groups in the KEGG nomenclature.

The Evolutionary Classification of Domains (ECOD) database <u>(Cheng et al. 2014; Cheng et</u> <u>al. 2015)</u> employs a hierarchical classification scheme that groups protein domains into nested tiers based on evolutionary relatedness (**Figure 1C**). In ECOD, closely related domains with highly similar sequences and structures are grouped into protein families (or F-groups). Related protein families are then collected into an X-group, the broadest evolutionary classification in ECOD, which emphasizes more distant evolutionary relationships. X-groups are hypothesized to have evolved from a single founder protein domain. We will refer to X-groups as *fold lineages* to differentiate this grouping from the narrower protein family grouping. There are 2,458 X-groups (fold lineages) and 15,144 F-groups (protein families) in ECOD. Linking fold lineages to KEGG orthologous groups can be done using the ECOD-curated database of hidden Markov models (HMMs). A detailed description of the procedure used to associate biochemical reactions, cofactors, enzyme orthology groups, and fold lineages is provided in the **Supplemental Methods**.

### Protein structure classification

Each fold lineage (X-group) in ECOD belongs to one of six structure classes based on the organization of ɑ-helices and β-sheets within the protein domain. ɑ- and β-proteins are comprised predominantly of ɑ-helices or β-sheets, respectively; ɑ+β proteins include both ɑ and β elements but they are structurally segregated from each other; ɑ/β proteins also contain both ɑ and β elements but, in this case, the two elements alternate along the chain; mixed proteins have features of both ɑ+β and ɑ/β proteins; and finally, “other” proteins lack well defined secondary structure elements. **Table S1** lists the structure classification of each fold lineage (X-group), based on the class–architecture mapping in **Table S2**.

### Standard metabolic network expansion

To model the stepwise evolution of biosphere-scale metabolism, the network expansion algorithm <u>(Handorf et al. 2005; Goldford et al. 2017)</u> was employed, which is described briefly here: First, a set of precursors, called *seed compounds*, are assumed. These seed compounds then undergo all possible reactions within a pre-defined set of allowed reactions, subject to reactant and cofactor requirements. The products of these reactions are then added to the pool of available compounds used in the next iteration. This process is repeated until no new compounds are discovered. Once a compound is produced within the network, it is never removed. Network expansion concerns reachability; it is not a kinetic model. The final set of compounds reached is referred to as the scope. *Ablation analyses* remove a component of the metabolic model — either a metabolite, a reaction, a fold lineage, or sets thereof — to assess its role in the expanding network. To ablate a single fold lineage, all reactions that require that fold lineage are removed. Likewise, ablation of a compound removes all reactions involving that compound, either as a reactant or a product. The network expansion implementation used in this work is available as a python package on GitHub (https://github.com/jgoldford/networkExpansionPy).

### Metabolic model

The reactions in our metabolic model were curated from KEGG <u>(Kanehisa and</u> <u>Goto 2000)</u> with added cofactor requirements <u>(Goldford et al. 2024)</u>. Some forward or reverse reactions were removed from the network based on thermodynamic constraints calculated by the component contribution method <u>(Noor et al. 2013)</u> using the eQuilibrator python API <u>(Beber et al,</u> <u>2022)</u>. If a reaction’s standard molar free energy is ever negative at pH = 7.0, pMg = 3.0, an ionic strength of 0.25 M, temperature T = 298.15, and allowing for a concentration range of 10^−7^–10^−1^ M for all reactants and products, the reaction was permitted. In the standard KEGG database, purines are required for purine synthesis. This circular causality blocks metabolic expansion early in the trajectory unless a non-autocatalytic pathway for purine synthesis is included <u>(Goldford et</u> <u>al. 2024)</u>. Thus, non-autocatalytic purine synthesis reactions were added to the model. These reactions were not given KO group or enzyme associations as none are known. Reactions with elemental inconsistency and reactions producing H_2_O_2_ before the discovery of molecular oxygen were removed. Allowed reactions are listed in **Table S3** and removed reactions are listed in **Table S4**.

### Seed set

Network expansion requires a set of seeding compounds from which the metabolic trajectory starts. The base seed set used in this work comprises 80 compounds including metals, inorganic molecules, CO_2_, H_2_S, H_2_, orthophosphate, ammonia, and 20 organic substrates that can be produced abiotically when reacting glyoxylate with pyruvate <u>(Muchowska et al. 2019)</u>. Many of the organic substrates are fundamental carbon compounds found in the reductive TCA cycle <u>(Braakman and Smith 2013)</u>. The base seed set also includes 10 amino acids with plausible prebiotic availability (glycine, *L*-alanine, *L*-aspartate, *L*-glutamate, *L*-isoleucine, *L*-leucine, *L*-proline, *L*-serine, *L*-threonine, *L*-valine) <u>(Higgs and Pudritz 2009; Longo and Blaber 2012)</u>. These amino acids are predicted to have been part of a primitive genetic code <u>(Copley et al. 2005)</u> and are sufficient to encoding stable, foldable proteins <u>(Akanuma et al. 2002; Longo et al. 2013; Yagi</u> <u>et al. 2021; Giacobelli et al. 2025)</u>. Nine additional seed sets, each with successively more compounds, were considered for the enzyme-gated network expansion analysis: Eight expanded seed sets, which correspond to the points of emergence of various cofactors (PLP, ATP, NADH, SAM, CoA, FAD, ThDP, and cobalamin) in the standard network expansion trajectory from the base seed set (and thus form nested sets); and one seed set that contains all metabolites in our metabolic model. Note that the ATP-expanded seed set also contains the canonical amino acids *L*-asparagine, *L*-cysteine, and *L*-glutamine, as well as the basic amino acid *L*-diaminobutyrate, which is hypothesized to have been part of the primitive genetic code <u>(Copley et al. 2005)</u>. The seed sets are listed in **Table S5**.

### Enzyme-gated network expansion

An implementation of enzyme-gated network expansion in Python is available at https://github.com/jgoldford/networkExpansionPy. A natural language description of the algorithm is presented in the main text. A pseudocode implementation of the algorithm is presented in **Figure S1**.

### Phyletic distribution score

Genome data were obtained from Genome Taxonomy Database (GTDB) <u>(Parks et al. 2022)</u> for archaea and bacteria (version 207) and EukProt (version 3) <u>(Richter</u> <u>et al. 2022)</u> for eukaryotes. HMM profiles from ECOD (version 279) <u>(Cheng et al. 2014)</u> were queried against the genomes using hmmsearch <u>(Mistry et al. 2013)</u> with the search space set to 106,052,079 sequences. Only hits with an independent E-value of less than 1 × 10^−5^ and an HMM coverage of greater than 60% were kept as potential hits. In cases where there was an overlap of more than 25% between two or more hits, only the hit with lower E-value was retained. Hits were assigned to X-groups (fold lineages) only when unambiguous association was possible. For each fold lineage, a phyletic distribution score, D, was calculated as:

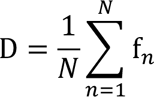

where f_n_ is the fraction of genomes containing the fold lineage within each phylum and N is the number of phyla. Phyletic distribution score was calculated separately for archaea, bacteria, and eukaryotes. Prokaryotic distribution score is the average of the archaea and bacteria phyletic distribution scores. Distribution scores for each fold lineage are provided in **Table S1**.

### Bacterial physiology annotation

Bacterial oxygen physiology annotations were obtained from a study that aggregated annotations from the Joint Genome Institute (JGI) Genomes Online Database (GOLD) <u>(Madin et al. 2020; Goldford et al. 2023)</u>. Taxa labeled “obligate aerobe” or “aerobe” were classified as aerobes, and taxa labeled “obligate anaerobe” or “anaerobe” were classified as anaerobes. NCBI taxonomy IDs were mapped to GTDB genome IDs using metadata provided on the FTP server from GTDB <u>(Parks et al. 2022)</u>. See **Table S6** for the full list of physiology annotations.

### Data and code availability

The analyses presented in this study are based on data retrieved from the KEGG database which can be accessed through www.kegg.jp, subject to their licensing policies and agreements. Processed datasets and code used in this study are available in **Supplementary Tables 1-15** and via a Zenodo repository (https://doi.org/10.5281/zenodo.15580734) and a GitHub repository (https://github.com/tseamuscorlett/enzyme-gated-network-expansion).

### Statistical analysis

Pearson correlations, Spearman’s correlations, and two-sided, non-parametric Mann–Whitney U tests were performed using the pearsonr, spearmanr and mannwhitneyu functions, respectively, from the scipy.stats python module, version 1.14.1 in python 3.12.4.

## Results and Discussion

### Enzymatically versatile ɑ/β proteins are key mediators of metabolism

Approximately 75% of the 7,678 reactions in our metabolic model are associated with one or more fold lineages (**Table S7**). However, just 396 fold lineages — less than 17% of those identified in the ECOD database — are represented. Nearly 90% of these “metabolic fold lineages” are detected in both archaeal and bacterial genomes, compared to just 42% for the full set of fold lineages (**Figure 2A, Figure S2, Table S1**). All protein structure classes (*e.g.*, α, β, α+β, α/β; see **Methods** for a description of each structure class) are present among the metabolic fold lineages, though ɑ/β structures are enriched 3.2-fold in the metabolic fold lineages relative to all fold lineages (**Figure 2B, Figure S2**). Likewise, the majority (60%) of enzyme orthology groups that catalyze a reaction in our metabolic model are associated with at least one protein domain that adopts an ɑ/β structure (**Figure 2C**). The number of reactions associated with each fold lineage (**Figure 2D**, y-axis) reveals a minority of exceptionally versatile fold lineages, with the three most versatile ones — Rossmann, TIM barrel, and flavodoxin — each adopting an α/β structure. The outsized role of ɑ/β proteins in metabolism has been noted previously <u>(Caetano-Anollés et al. 2007; Ma et al. 2008)</u>, and may relate to an intrinsic preference for, or a historical association with, (di)nucleotide cofactors (<u>Longo et al. 2020; Yanai et al. 2025</u>).

**Figure 2.**
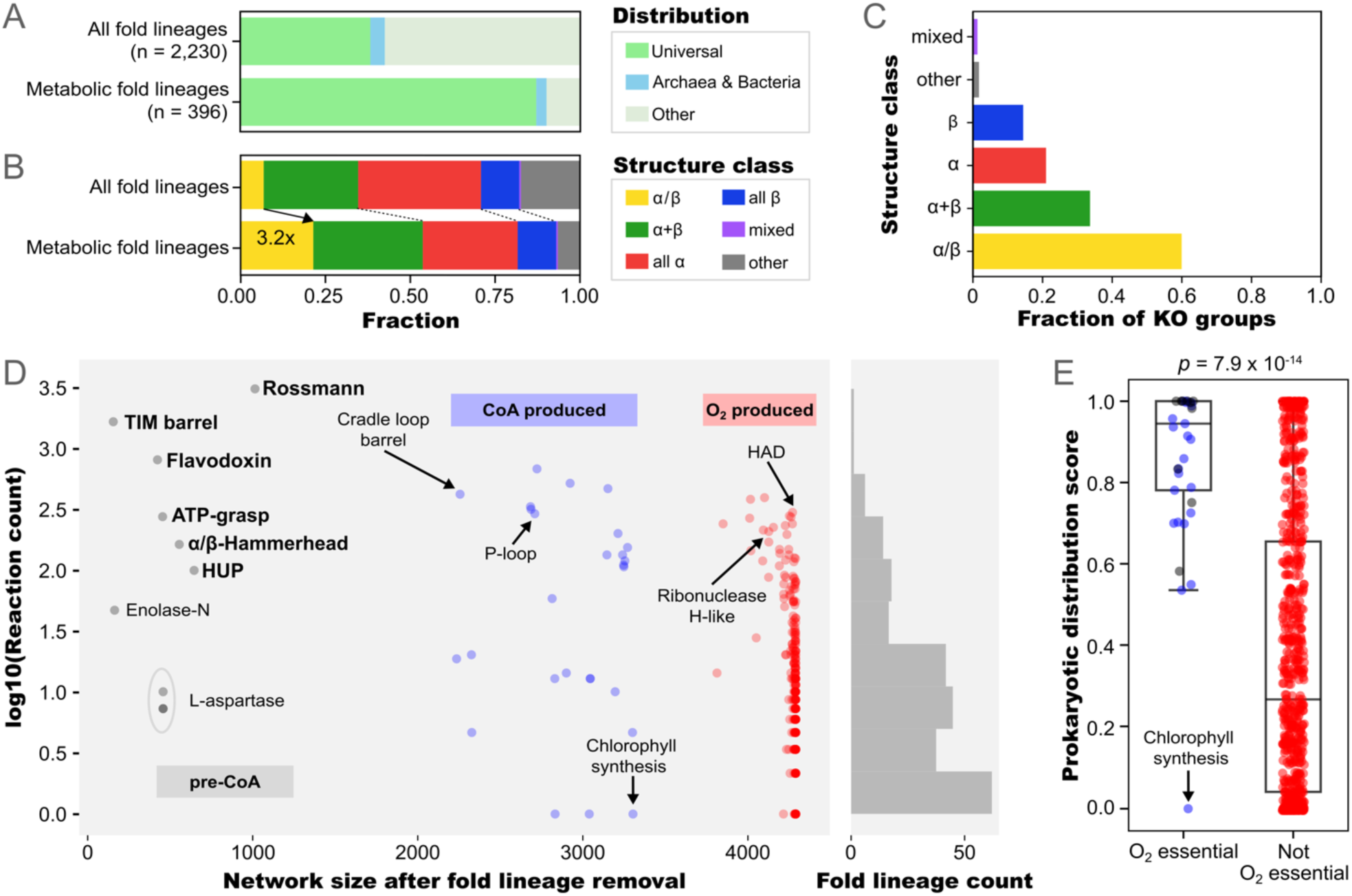
The metabolic fold lineages. **A**. Distribution of all fold lineages in ECOD and the subset of metabolic fold lineages. Universal refers to fold lineages present in all three domains of life: archaea, bacteria, and eukaryotes. **B**. Structure class of all fold lineages in ECOD and the subset of metabolic fold lineages. **C**. The fraction of enzyme orthologous groups associated with metabolic reactions in our model that contain fold lineages from each structure class. **D**. The number of associated reactions (y-axis) and the network size from standard network expansion after single fold lineage ablation (x-axis) for each metabolic fold lineage. CoA- (blue) and O_2_-containing (red) networks are indicated. **E**. Prokaryotic distribution scores (see **Methods** for definition) of fold lineages that are essential for reaching O_2_ versus those that are not, based on the single fold lineage ablation analysis in panel C.

### Both versatile and specialist fold lineages enable metabolic innovation

Counting the number of reactions associated with a fold lineage emphasizes the role of versatility. Some fold lineages, however, may catalyze a small number of essential reactions. To account for these cases, we employed metabolic network expansion <u>(Handorf et al. 2005)</u>.

Metabolic network expansion predicts the relative discovery order for a set of reactions based on their substrate requirements. Given a set of “seed compounds” that are provided at the beginning of the simulation, reachable reactions are activated to generate new metabolites that, in turn, activate new reactions. The process is repeated until no new metabolites can be discovered — either because all metabolites have been discovered or because a key reactant or cofactor is absent and further expansion is not possible. The set of all discovered metabolites is referred to as the scope. By removing the reactions that are dependent on a fold lineage (“fold lineage ablation”) and calculating the resulting scope, specialist enzymes that catalyze a small number of essential reactions can be uncovered.

Using seed compounds based on a primitive Wood-Ljungdahl pathway <u>(Muchowska et al. 2019)</u> and TCA cycle <u>(Braakman and Smith 2013)</u> (see **Methods** for more details), network expansion was performed after the independent ablation of each fold lineage (x-axis of **Figure 2D** and **Table S8**). Foremost, the ablation analysis highlights the hierarchical structure of metabolism <u>(Raymond and Segrè 2006)</u>, with the resulting networks forming nested layers: Small networks that lack both CoA and O_2_, medium-sized networks that include CoA but lack O_2_, and large networks that include both CoA and O_2_ (**Figure 2D**). As expected, both versatile fold lineages (*e.g.*, Rossmann, TIM barrel, and flavodoxin) and specialist fold lineages (*e.g.*, the three *L*-aspartase fold lineages that mediate purine biosynthesis and X-group 3997, which mediates chlorophyll biosynthesis) were essential for complete metabolic expansion (**Figure 2D**). To our surprise, however, ablation of several versatile fold lineages (*e.g.*, the haloacid dehydrogenase lineage, abbreviated HAD) had only a minor impact on the scope. In total, only 37 fold lineages resulted in a contraction of the scope by >20% upon individual ablation. The fact that these 37 fold lineages are more widely distributed across prokaryotes than other fold lineages (*p* = 7.9×10^-14^, Mann-Whitney U test; **Figures 2E**), and include both versatile and specialist fold lineages, highlights the key contributions that both kinds of fold lineages make to metabolic innovation.

### Biosphere-scale metabolism is rich with alternatives

Does the core of metabolism require just 37 fold lineages, with all others playing a supporting role? Ablation of all fold lineages except the 37 that significantly contract the scope upon individual ablation (*i.e.*, network expansion with just 37 fold lineages) resulted in a network with just 290 metabolites (∼7% of full scope). The robustness we observe to single fold lineage ablations, but not to this multi-fold lineage ablation, is due to two main factors: First, approximately 12% of enzyme-catalyzed reactions can be catalyzed by entirely distinct sets of fold lineages (**Figure S3**). Second, the same metabolite can be achieved by alternative pathways, reflecting the composite, biosphere-scale nature of our metabolic model. Thus, ablating a single fold lineage (or a single reaction <u>(Goldford et al. 2024)</u>) reveals the first point along the expansion trajectory where it becomes essential for further network growth despite these two forms of redundancy.

Neither an analysis of enzyme versatility (**Figure 2D**, y-axis) nor of enzyme essentiality upon ablation (**Figure 2D**, x-axis) has provided a satisfying readout of the co-evolution of metabolism and enzymes, as both approaches emphasize the roles of just a small subset of fold lineages. A model that explicitly accounts for the co-evolution of metabolism and fold lineages, as well as the interplay between specialist and versatile fold lineages, is required. It is unclear, however, if a reachability analysis in the spirit of standard network expansion can be made suitable for this task: Is a stepwise trajectory constrained by fold/metabolism co-evolution able to meaningfully inform the emergence order of individual fold lineages? Or will the complex web of relationships between metabolites, reactions, cofactors, and enzymes preclude efforts to constrain co-evolutionary histories in the absence of external information such as phylogenetics?

### Enzyme-gated network expansion: A model for the joint evolution of metabolism and enzymes

We propose an algorithm to reconstruct the relative order of fold lineage emergence within the context of a growing metabolic network using an extended network expansion approach. We refer to this algorithm as *enzyme-gated network expansion* (**Figure 3A**) to emphasize the constraining role played by fold lineages.

**Figure 3.**
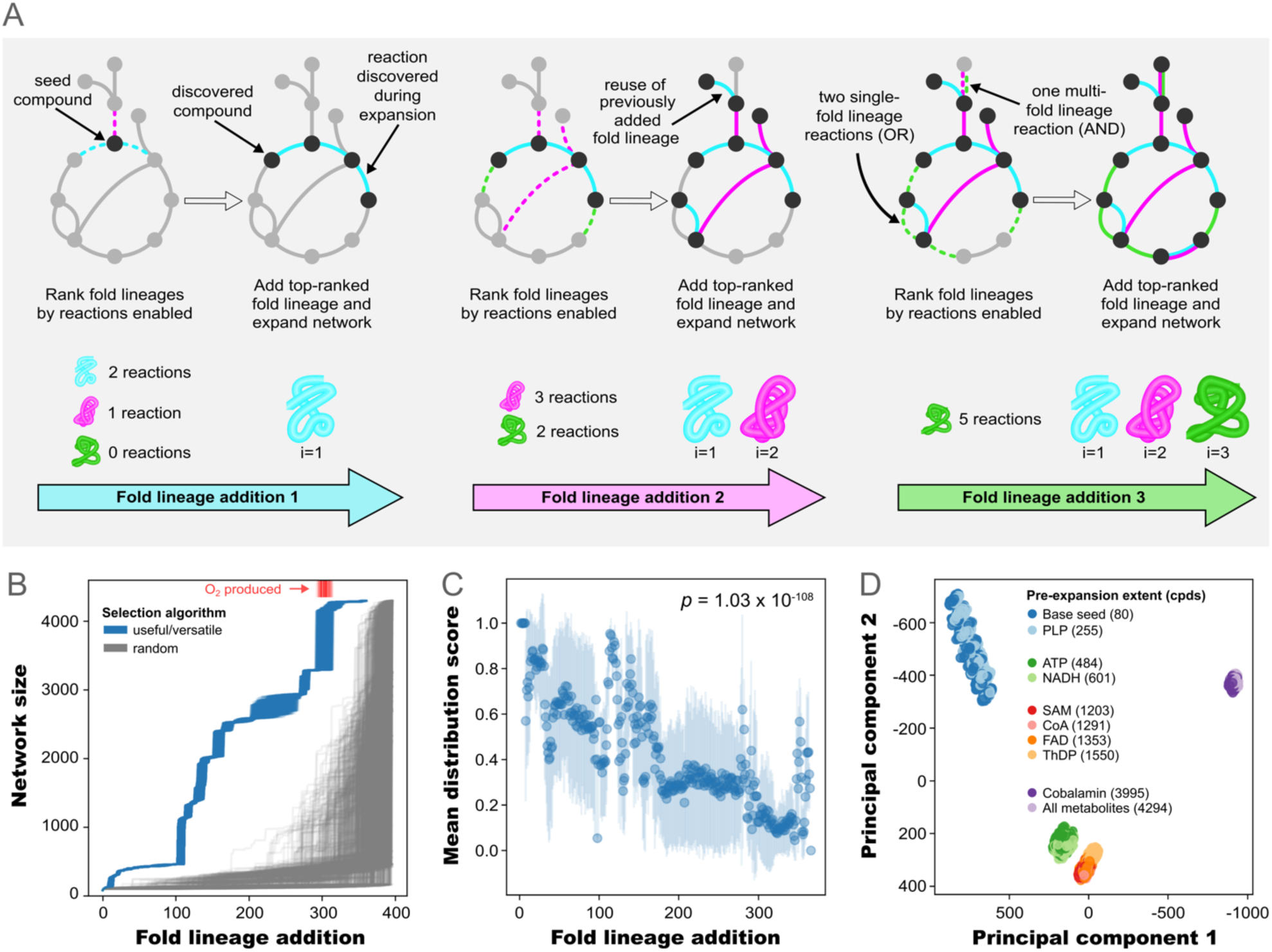
Enzyme-gated network expansion. **A**. The enzyme-gated network expansion algorithm. See main text and **Methods** for additional details. **B**. Metabolic growth trajectories (network size) when selecting the most useful/versatile fold lineage (blue) or a random fold lineage (grey) for 1,000 runs each. The red rug plot at the top indicates the iteration where molecular oxygen is discovered in the useful/versatile runs. **C**. Average prokaryote distribution score of fold lineages across 1,000 runs as a function of fold lineage addition order. Error bars are standard deviations. Indicated p-value calculated from the Spearman’s rank correlation. Fold lineage additions for panels B and C are based on the base seed set of compounds. **D**. Principal component analysis of the fold lineage addition orders from all trajectories across all ten seed sets (1,000 independent runs per seed set, 10,000 runs in total). The compounds for each seed set are listed in **Table S5**. Panels B and C for each seed set are provided in **Figures S8** and **S9**.

We model the evolution of reactions and fold lineages as a stepwise process with discrete emergence events. Here, new reactions can only be discovered if the requisite set of fold lineages, reactant compounds, and cofactors are available. However, unlike metabolites and cofactors, fold lineages are not endogenously generated within the reaction network, but must be introduced externally. Thus, each simulation begins with a set of seed compounds and no fold lineages. At first, only enzyme-independent reactions are active, which are insufficient for significant metabolic expansion (yielding just 2% of the scope). To activate more reactions, fold lineages are added in an independent, sequential fashion. After each fold lineage addition, all reactions that can be catalyzed by that fold lineage — alone or in combination with already discovered fold lineages — are activated and network expansion proceeds until no new metabolites can be reached. By allowing as much network expansion as possible before the addition of the next fold lineage, we assume that the discovery of new fold lineages is infrequent compared to their functional diversification. The process is repeated until no new reactions can be activated.

The order of fold lineage addition is selected based on a combination of utility and versatility in the context of the growing network. For a fold lineage to be added at a given point in the trajectory, it must be able to catalyze at least one reaction given the current set of metabolites and fold lineages (utility requiring). Note that the newly catalyzed reaction(s) do not need to produce new metabolites, since (a) new reactions between previously discovered metabolites, (b) previously discovered reactions catalyzed by a different fold or set of folds, and (c) entirely new reactions are treated as equivalent when selecting a fold lineage. If multiple fold lineages meet this criterion, the fold lineage that activates the most reactions is selected (versatility favoring).

This second selection criterion stems from the intuition that domain emergence can be supported through interactions with a cognate ligand <u>(Alva et al. 2015; Yagi and Tagami 2024)</u>. Thus, the emergence of a fold lineage that can interact with a large pool of cognate ligands (versatile) may be more likely than the emergence of a specialist fold lineage with only a small number of cognate ligands. In the event of a tie, selection is random among the most versatile fold lineages. Consequently, we describe an envelope of possible trajectories given these simple selection criteria.

### Fold lineage emergence order is well constrained by enzyme-gated network expansion and comports with phyletic distribution

Using the enzyme-gated network expansion approach, we attempted to reconstruct the co-evolutionary history of fold lineages and metabolism. However, when starting from the base seed set, 93 sequential fold lineage additions yielded only 455 compounds before the simulation terminated, indicating that one or several essential reactions could not be activated. An analysis of fold lineage-reaction associations revealed that nearly 50% of fold lineages cannot operate without a partner fold lineage. As a result, the growing network reaches a point where any single fold lineage addition does not lead to the discovery of a new reaction with this minimal algorithm. To overcome this challenge, reactions that can be catalyzed by two or more fold lineages are considered for activation by the simultaneous addition of the smallest possible set of fold lineages (usually pairs). As with single fold lineages, versatility is favored and fold set selection is random if multiple fold sets are possible. With this adjustment, the full set of metabolites in our metabolic model is achieved (**Figure 3B**). A pseudocode implementation of the enzyme-gated network expansion algorithm is presented in **Figure S1**.

The majority of fold lineages (79%, averaged over 1,000 simulation runs) — including both versatile and specialist fold lineages — are required for expansion to an O_2_-containing metabolic network with enzyme gating (**Figure 3B**). Plotting the results of 1,000 independent runs reveals a surprisingly narrow distribution of metabolic trajectories (**Figure 3B**, useful/versatile). These trajectories are highly similar to that of the standard network expansion (**Figures S4** and **S5**; average Spearman’s rank correlation coefficient of 0.85), and enzyme gating does not significantly alter the patterns of pathway emergence (**Figure S6A**, **Table S9**) or the order of carbon fixation pathway discovery (**Figure S6B, Table S10**) relative to standard network expansion.

Does enzyme-gated network expansion meaningfully constrain the relative emergence order of fold lineages? Several pieces of evidence suggest that it does: First, the standard deviation of the addition step is small (less than 20 fold lineage addition steps) for 96% of fold lineages (**Figure S7**). Second, fold lineages added in a random order — with no requirement for activating a reaction and no multi-fold lineage additions — can serve as a point of comparison to the useful/versatility selection criteria (**Figure 3B**). Whereas random fold addition orders yield significant growth only at the end of the trajectory, application of the useful/versatility selection criteria maintains modest growth of the network throughout the trajectory — though bursts in expansion are still apparent. Finally, the fold lineage sampling along the trajectory is highly correlated with the phyletic distribution of fold lineages in prokaryotes (**Figure 3C**). These results suggest that the predicted order of fold lineage addition by enzyme-gated network expansion meaningfully reflects aspects of the relative ordering of their emergence as enzymes.

### At what point in the history of metabolism did enzymes emerge?

The earliest metabolic reactions were likely catalyzed by minerals <u>(Muchowska et al. 2019)</u> and cofactors <u>(Goldman and Kacar 2021)</u> prior to the emergence of enzymes. At what level of metabolic complexity did fold lineage emergence begin, and can our approach reveal signatures of this transition? To explore these questions, ten sets of seed compounds, ranging from 2% to 100% of metabolites in our model, were analyzed using enzyme-gated network expansion (**Figures 3D** and **S8**). The seed sets of intermediate size correspond to the inventory of metabolites available at the point of discovery of eight cofactors in the standard network expansion model: pyridoxal phosphate (PLP, 6% of metabolites), adenosine triphosphate (ATP, 11%), nicotinamide adenine dinucleotide (NADH, 14%), S-adenosyl methionine (SAM, 28%), coenzyme A (CoA, 30%), flavin adenosine dinucleotide (FAD, 32%), thiamine diphosphate (ThDP, 36%), and cobalamin (93%).

To our surprise, a principal component analysis of the fold lineage orders (1,000 runs per seed set) revealed just three major clusters: a pre-nucleotide cofactor seed set, a post-nucleotide seed set (which breaks into two sub-clusters), and a post-O_2_ seed set. All seed sets constrain the relative order of fold lineage emergence (**Figure S8**) and have strong rank correlation between phyletic distribution and addition order (Spearman’s rank correlation, *p* < 1.3 x 10^-76^; **Figures 3C** and **S9**). However, with increasingly larger seed sets, fold lineage addition order more strongly reflects the number of reactions associated with a fold lineage (versatility-emphasizing), suppressing the role of specialist lineages in catalyzing essential reactions (**Figure S10**).

Various lines of evidence highlight the post-nucleotide seed set as the “natural” position for the enzymatic takeover of metabolism. Foremost, coded protein synthesis depends on nucleotides, meaning that pre-nucleotide synthesis of contemporary domains is unlikely. The observation that pre-nucleotide seed sets result in (a) long intervals of single fold lineage additions without significant metabolic expansion and (b) the requirement for concurrent addition of pairs or larger groups of fold lineages to discover new metabolic reactions early in the trajectory is consistent with a too-early initiation of enzyme gating (**Figure S11**). As enzymes likely predate the biological production of molecular oxygen <u>(Sousa et al. 2016)</u>, we conclude that the nucleotide-containing seed set is the most reasonable reflection of early enzyme history. Given the strong similarity between nucleotide-containing seed sets (**Figures 3D** and **S11**), we focus on the ATP-containing seed set going forward.

### The first metabolic fold lineages: TIM Barrel and the Rossmannoids

A joint history of metabolite and fold lineage discovery based on the ATP seed set is presented in **Figure 4A** (see **Tables S11** and **S12** for the discovery order of metabolites and fold lineages, respectively). Previously, metabolic evolution was observed to be bursty or punctuated, with periods of innovation interspersed with periods of quiescence <u>(Goldford et al. 2024)</u>. Enzyme gating did not fundamentally change this property of the substrate trajectories (see also **Figures 3B** and **S12**). It does, however, assign an emergence order to the earliest fold lineages, which have proven recalcitrant to relative dating by other methods due to their near-absolute conservation across the tree of life <u>(Winstanley et al. 2005; Edwards et al. 2013)</u>.

**Figure 4.**
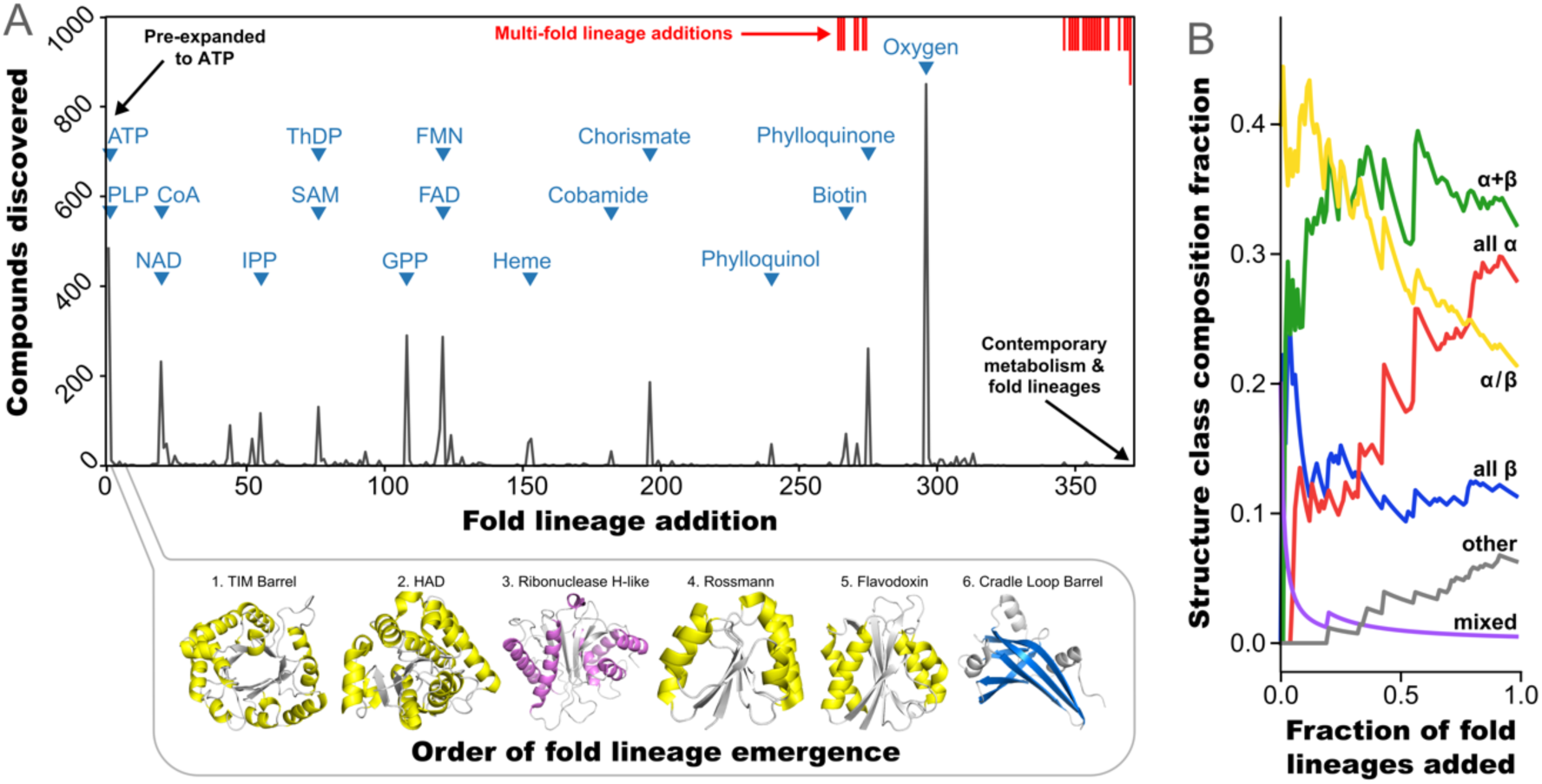
A coevolutionary history of fold lineages, metabolic reactions, and metabolites. **A**. Fastest growing trajectory (cumulative plot with the greatest integrated area) out of 1,000 runs for the ATP seed set. Select metabolites and the first 6 fold lineages, colored according to structure class, are indicated. The position of metabolite labels on the y-axis is arbitrary. The red rug plot along the top of the figure denotes fold lineage addition steps where two (short line) or three (long line) fold lineages had to be added simultaneously for a new reaction to be discovered. The average orders of emergence of all fold lineages across 1,000 runs for each seed sets are provided in **Table S13**. **B**. Average structure class composition along the trajectory for the ATP seed set. Other seed sets are provided in **Figure S14**. The “mixed” structure class refers to fold lineages with both ɑ/β and ɑ+β elements.

The first six fold lineages (**Figure 4A**) — which appear in the same order across all 1,000 independent ATP seed set runs, and are similarly positioned in other seed sets (**Figure S13, Table S13**) — are the TIM barrel, HAD, ribonuclease H-like, Rossmann, flavodoxin, and cradle-loop barrel. The profound metabolic importance of the TIM barrel is well established <u>(Goldman et al. 2016)</u>, as are its evolutionary relationships to flavodoxin <u>(Farías-Rico et al. 2014)</u> and other Rossmannoids (in this case, HAD and Rossmann) <u>(Kolodny et al. 2021; Qiu et al. 2022)</u>. The early emergence of HAD and ribonuclease H-like, which ablation studies classified as versatile but non-essential (**Figure 2D**), likely relates to the large repertoire of simple reactions that these fold lineages catalyze. That 4/6 of the earliest fold lineages belong to the α/β structure class and one adopts a “mixed” structure (Ribonuclease H-like), having significant α/β and α+β character, is consistent with hypotheses that α/β structures were the earliest entry point into folded domains based on phyletic distribution <u>(Caetano-Anollés et al. 2007)</u>, sequence-structure fragment sharing <u>(Nepomnyachiy et al. 2014)</u>, and patterns of protein structure similarity <u>(Choi and Kim</u> <u>2006)</u>. Moreover, TIM barrels, Rossmannoids, and some β-barrel folds adopt repetitive structures (*e.g.*, TIM barrel has a pseudo axis of *C_4_* symmetry and the Rossmann has a β-strand order of 3-2-1-4-5-6, suggestive of an early duplication event) underscoring symmetry and repetition as organizing principles in early protein evolution <u>(Eck and Dayhoff 1966)</u>. We have previously hypothesized that the early dominance of the α/β structure class likely relates to facile emergence and functional potential under the constraints of a limited amino acid alphabet and short peptide length <u>(Romero Romero et al. 2018; Longo et al. 2020; Longo et al. 2020; Vyas et al.</u> <u>2021)</u>. However, we observe a dilution of α/β fold lineages as the trajectory proceeds (**Figure 4B**) — with α and α+β fold lineages being preferred later in the trajectory. This result may suggest that α/β fold lineages are not fundamentally more evolvable or functionally capable under present conditions, or that other structure classes may be uniquely suited to perform emergent metabolic roles.

Although β-proteins play a more limited role in metabolism (**Figures 2B**, **2C,** and **4B**), remarkably, the first β-protein predicted to have emerged by enzyme-gated network expansion is the cradle-loop barrel. This fold lineage plays essential roles in information processing and encompasses several small, highly traversable β-barrel folds <u>(Yagi and Tagami 2024)</u>. Small β-folds have also been suggested to play a founder role, particularly among ribosomal proteins <u>(Petrov et al. 2015; Kovacs et al. 2017)</u>. The fact that a ribosome-centric view does not highlight α/β as being ancient may reflect different emergence filters for ribosome-associated and metabolism-associated folds — for example, nucleic acid binding versus independent foldability and nucleotide binding, respectively.

### Bursts of metabolic innovation recruit fold lineages young and old

When new metabolites arise in the biosphere, they can drive the emergence of new fold lineages or be acted upon by pre-existing fold lineages performing a new reaction (*reuse*). The history of fold lineage reuse upon the emergence of new metabolites for a representative ATP seed set run is presented in **Figure 5A**. While approximately half of fold lineages are used when they are introduced and never again, 54% (216/396) of lineages are reused at multiple points along the trajectory (**Figure S15**). Some lineages, particularly the early-emerging, versatile fold lineages, exhibit near constant reuse along the trajectory. The Rossmann lineage, for example, is reused after ∼30% of fold lineage additions, likely due to its close association with various cofactors and diverse repertoire of cooperating domains. Other lineages lie dormant for long periods before finding a new use; X-group 211, for example, is the 99^th^ fold lineage added and performs carbon-sulfur lyase chemistry. This fold lineage is then later reused to catalyze a reaction involving O_2_ upon addition of the 296^th^ fold lineage. About 70% (152/216) of the fold lineages with reuse catalyze a reaction with a new 2-digit EC (Enzyme Commission) number sometime later, usually within 30 fold lineage additions of their emergence (**Figure S16**). Peaks in fold lineage reuse (**Figure 5A**, upper) correspond to significant events of metabolic growth, and late peaks increase in size as more established fold lineages are available to draw from. The number of required fold lineages per discovered reaction modestly increases over the expansion (**Figure S17**) as key lineages accrete cooperating domains (see **Figure S18** for an analysis of the TIM barrel).

**Figure 5.**
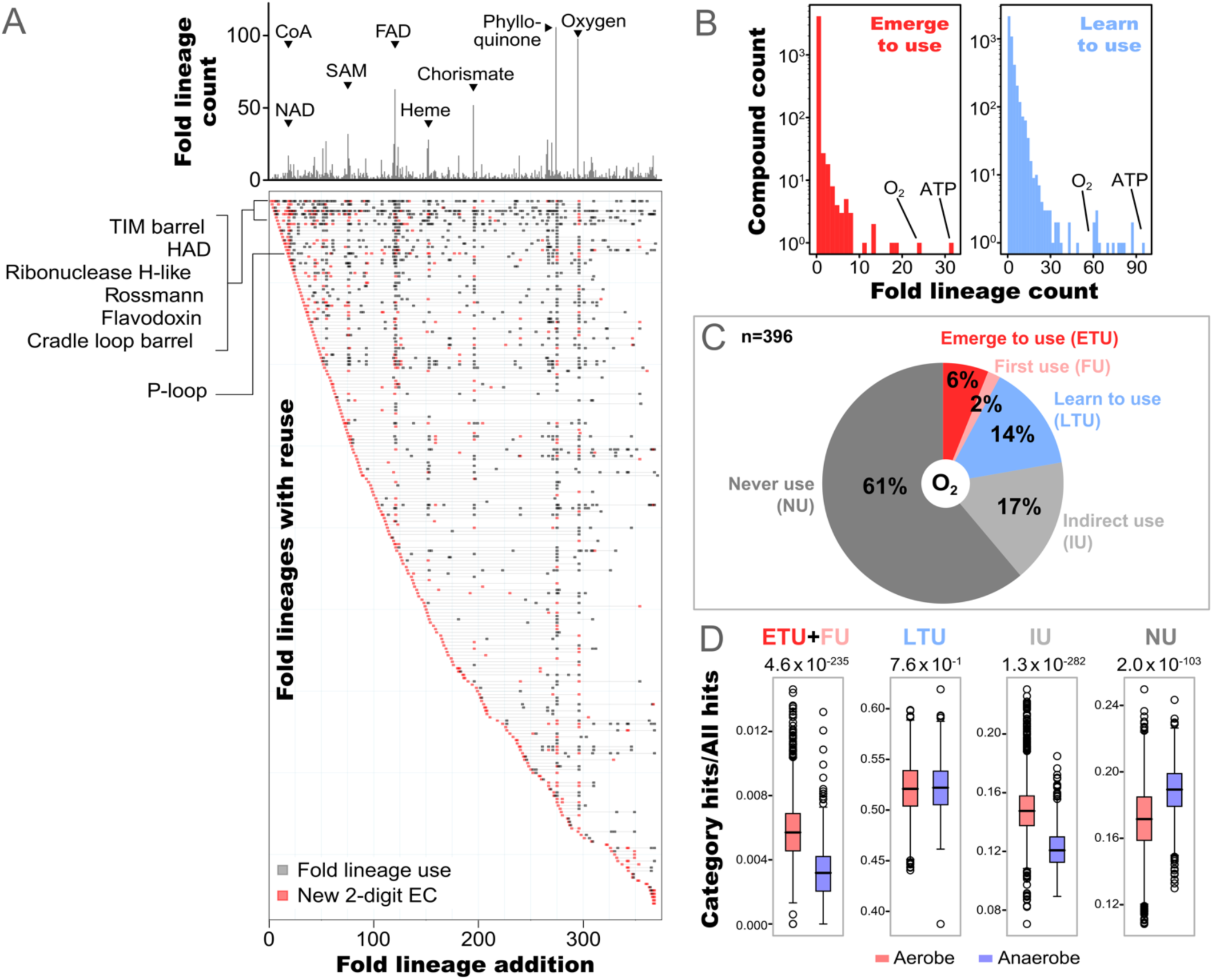
Emergence and reuse of fold lineages. **A**. A global view of fold lineage reuse calculated from the trajectory in Figure 4A. Total fold lineage use counts at various points in the expansion (upper) and the patterns of fold lineage emergence and reuse along the trajectory (lower). Fold lineages are shown one per row along the y-axis, while the x-axis indicates points along the trajectory where a fold lineage is associated with a new reaction. The fold lineages have been ordered from top to bottom by their order of emergence. Only fold lineages that catalyze a reaction at 2 or more points in the trajectory are shown. Points where a fold lineage catalyzes a reaction with a new 2-digit EC number are indicated in red. **B**. Histogram of the number of fold lineages that emerge to use (left) or learn to use (right) a given compound. ATP and molecular oxygen are both strong drivers of fold lineage emergence and reuse. **C**. Breakdown of fold lineage relationships with oxygen. See main text for description of each class. Panels B and C are calculated from 1,000 runs of the ATP seed set. Only classifications consistent across ≥75% of runs are assigned. In the case of oxygen, all fold lineages satisfy this threshold. A full list of metabolic fold lineages and their category is available in **Table S14**. **D**. Enrichment of fold lineages associated with each oxygen-use category in bacteria with predicted aerobic or anaerobic physiologies. For each genome (1,630 aerobes and 1,099 anaerobes), the occurrence of fold lineages within a given usage category was divided by the occurrence of all ECOD fold lineages. Indicated p-values were calculated by Mann-Whitney U tests.

### Reuse outpaces emergence: The case of molecular oxygen

To quantify the impact of metabolite discovery on the protein universe, each fold lineage was grouped into one of five classifications with respect to a given metabolite (see **Supplementary Methods** and **Figure S19**): Fold lineages whose first reactions all require the metabolite (*emerge to use*, ETU); fold lineages for which some of their first reactions require the metabolite (*first use*, FU); fold lineages that catalyze reactions involving the metabolite sometime after their emergence (*learn to use*, LTU); fold lineages that catalyze reactions that are made possible by the metabolite, but do not catalyze reactions with the metabolite directly (*indirect use*, IU); and fold lineages that are not associated with the metabolite in any way (*never use*, NU). In cases where multiple classifications apply, the order of precedence was ETU/FU>LTU>IU.

We find that molecular oxygen is one of the strongest drivers of both fold lineage emergence and reuse (**Figure 5B**). The emergence of oxygenic photosynthesis occurred approximately 3 billion years ago <u>(Crowe et al. 2013)</u> and added a layer of O_2_-associated reactions around a largely anaerobic core, a metabolic organization referred to as hierarchical modularity <u>(Raymond and Segrè 2006)</u> (**Figure 2D**). The modest rewiring of interior metabolic layers that resulted from the advent of O_2_, however, stands in stark contrast to its consequence within the protein universe, where many pre-existing fold lineages acquired new functions. Our model predicts that oxygen utilization was a primal function for 8% of metabolic fold lineages (ETU or FU; **Figure 5C**, **Table S14**) whereas almost two times more fold lineages (14%) acquired the ability to use oxygen later in the trajectory, sometime after their emergence as enzymes (LTU). 65% of versatile fold lineages (having ≥100 associated reactions) fall into the LTU category (**Figure S20**). We note, however, that while the consequences of biological O_2_ production had sweeping impacts on established fold lineage functions, it was only possible due to the existence of several specialist fold lineages, such as those responsible for synthesizing chlorophyll.

To further test the predicted fold lineage-oxygen emergence relationships, we analyzed the domain composition of aerobic and anaerobic bacteria <u>(Madin et al. 2020; Goldford et al. 2023)</u> (**Table S6**). We find that category-wise fold lineage enrichments (**Figure 5D**) are fully consistent with expectation: ETU+FU and IU fold lineages are enriched in aerobes. NU fold lineages, on the other hand, are more enriched in anaerobes. Finally, LTU fold lineages, which presumably emerged before oxygen and only later catalyzed an oxygen-associated reaction, are equally common in both anaerobes and aerobes. Thus, signatures of fold lineage adaptation to oxygen predicted by enzyme-gated network expansion are manifested as distinct distribution patterns across bacteria with different physiologies, suggesting that our approach can approximate the relative order of enzyme functions.

### Limitations and outlook for models of metabolic evolution with strict causality

The present model enforces strict enzyme-reaction and cofactor-reaction dependencies and does not consider the kinetic constraints that must be satisfied for completing metabolic pathways. These decisions simplify implementation and interpretability but can introduce errors in cases where the essential source data (*e.g.*, biochemical reactions) are missing or incorrect, as was the case for non-autocatalytic purine synthesis <u>(Goldford et al. 2024)</u>. The strengths and limitations of the network expansion algorithm have been discussed in detail <u>(Goldford and Segrè</u> <u>2018)</u>, and apply to this approach as well. In addition, we note that this approach reflects the emergence of a fold lineage as a metabolic enzyme, and does not account for other types of functional roles that may have emerged earlier. Future work can begin to integrate alternative historical records, such as by incorporating homology relationships *between* fold lineages <u>(Farías-Rico et al. 2014; Longo et al. 2020; Kolodny et al. 2021)</u>, or by developing frameworks with relaxed causality through careful coarse graining.

## Concluding Remarks

We introduce a new approach to interpret the record of enzyme evolution embedded within the graph structure and reaction–fold lineage associations of biosphere-scale metabolism.

Remarkably, we recover a sequential ordering of fold lineage emergence in which each new fold lineage was largely sufficient on its own to drive metabolic expansion. This record of protein history is independent from (though found to be correlated with) phyletic distribution, and can provide evidence for the emergence order of ancient fold lineages that reflects both fold lineage utility and essentiality. The first metabolic fold lineages were likely a group of related ɑ/β lineages, but the dominance of the ɑ/β class was diluted over time. Differences between metabolic-centering and ribosome-centering perspectives — particularly with respect to the emergence order of protein classes — may provide clues for multiple fold origins. In addition, we provide a snapshot of how the protein universe responds to metabolic upheaval, with preference for reuse of pre-existing fold lineages over the emergence of entirely new fold lineages. This work adds a new archive to the goal of synthesizing diverse historical records to reconstruct the earliest stages of protein evolution.

## Acknowledgements

Shawn E. McGlynn helped manually annotate KEGG orthology group interdependence. This study was supported by the Human Frontier Science Program Grant Number RGEC29/2025 (LML) with the award DOI https://doi.org/10.52044/HFSP.RGEC292025.pc.gr.230124 and the National Aeronautics and Space Administration (NASA) award number 80NSSC25K7873 (LML) and 80NSSC23K1357 (JG).

## Author Contributions

Conceptualization: TC, HS, JG, LML

Methodology: TC, HS, JG, LML

Software: TC, HS, JG

Validation: TC, HS

Formal Analysis: TC, HS, LML

Investigation: TC, HS

Resources: HS, JG, LML

Data curation: TC, HS, JG, LML

Writing — Original Draft: TC, ES, LML

Writing — Review & Editing: HS, JG

Visualization: TC, LML

Supervision: ES, HS, JG, LML

Project administration: LML

Funding acquisition: LML

