## Supplementary Figures for "The history of enzyme evolution embedded in metabolism"

### Table of Contents

**Supplemental Methods** ------------------------------------------------------------------------------ 3
**Figure S1.** Pseudocode of the enzyme-gated network expansion algorithm ---------------- 5
**Figure S2.** Phyletic distribution of fold lineages -------------------------------------------------- 7
**Figure S3.** Mapping of reactions to fold lineage sets -------------------------------------------- 8
**Figure S4.** Compound discovery order: enzyme-gated vs. standard network expansion--- 9
**Figure S5.** Standard deviation of compound discovery orders -------------------------------- 10
**Figure S6.** Pathway emergence ----------------------------------------------------------------------11
**Figure S7.** Standard deviation of fold lineage addition orders --------------------------------- 13
**Figure S8.** Network expansion trajectories with seed set pre-expansion --------------------14
**Figure S9.** Average prokaryotic distribution at each fold lineage addition step --------------15
**Figure S10.** Trends in seed set pre-expansion and fold lineage selection -------------------- 16
**Figure S11.** Periods of multi-fold lineage discovery ---------------------------------------------18
**Figure S12.** Burst analysis ------------------------------------------------------------------------- 19
**Figure S13.** Comparison of mean fold lineage addition orders --------------------------------20
**Figure S14.** Protein structure class discovery --------------------------------------------------- 21
**Figure S15.** Fold lineage reuse ----------------------------------------------------------------------22
**Figure S16.** Fold lineage neofunctionalization -------------------------------------------------- 23
**Figure S17.** Fold lineage set complexity ---------------------------------------------------------- 23
**Figure S18.** Domain accretion history ------------------------------------------------------------ 24
**Figure S19.** Fold lineage-oxygen emergence relationships ----------------------------------- 25
**Figure S20.** Fractions of $O_2$ reactions in LTU and IU fold lineages ---------------------------- 26
**Supplemental References** --------------------------------------------------------------------- 27

#### Supplemental Methods

*Reaction-KO group associations*. Reaction-KO group associations were extracted from the KEGG database, accessed through the KEGG web interface or the KEGG REST API (either directly or through Biopython) (Kanehisa and Goto 2000; Cock et al. 2009). Three types of annotations were used: a) direct KO-reaction mappings, b) KO-reaction associations linked through KEGG modules, and c) KO-reaction associations linked though EC numbers. For reactions associated with exactly one KO group, no further curation was performed. For reactions that map to multiple KO groups, all KO groups were reviewed to annotate interdependence relationships between KO groups. KO groups were generally assumed to be isozymes capable of catalyzing the same reaction independently unless literature searching indicated that they are interdependent components (such as subunits of a protein complex). KO groups labeled as membrane anchors, effectors, coupling domains, or auxiliary domains were considered nonessential and not included as reaction dependencies.

*Fold lineage-KO group associations*. Associations between KEGG orthologous (KO) groups (Kanehisa and Goto 2000) and ECOD X-groups (Cheng et al. 2014) were constructed as follows: Genes from 8,030 KO groups were clustered at 80% sequence identity by CD-HIT (Fu et al. 2012) using a word size of 5. Hidden Markov models (HMMs) from version 279 of the ECOD database were queried against the representative KEGG genes of each KO group using hmmsearch (Mistry et al. 2013) with the search space set to 106,052,079 sequences. Domains with an independent E-value of less than $1 \times 10^{-4}$ and an HMM coverage of greater than 70% were considered potential hits. Hits were assigned to gene regions in order of increasing E-value using the “envelope coordinates.” If two hits overlapped by more than 30% of either hit, the hit with the higher E-value was discarded. Finally, reconciled hits associated with a gene were mapped to their corresponding X-groups when such mappings are unambiguous. X-group sets corresponding to at least 20% of the representative genes within a KO group were taken to be catalytically competent. Nested sets, in which one observed X-group set is a subset of another X-group set for a given KO group, were retained. Note that X-groups are referred to as fold lineages throughout the main text.

*Reaction-fold lineage associations*. Based on the fold lineage-KO group and reaction-KO group associations described above, reaction-fold lineage associations were constructed via their mutual linkage through KO groups.

*Reaction-cofactor associations*. Detailed cofactor dependencies for 5,259 reactions were manually reconciled from annotations in UniProt (UniProt Consortium 2023), Expasy

(Gasteiger et al. 2003), PDBe (Armstrong et al. 2020), EBI (Rodriguez-Tomé et al. 1996), and KEGG (Kanehisa and Goto 2000) (**Table S3**). Each cofactor is associated with one or several closely related KEGG compounds (**Table S15**) that can, to a reasonable approximation, satisfy the mechanistic role of that cofactor class. A full description of reaction-cofactor associations, and their implementation in the standard network expansion, was described previously (Goldford, 2024).

*Fold lineage-metabolite emergence relationships*. Each fold lineage-metabolite pair was classified into one of five emergence relationships. The classification proceeded as follows: For each fold lineage, its *first reactions* — reactions catalyzed by the fold lineage in the first network expansion step immediately after addition — were identified across 1,000 runs of enzyme-gated network expansion with the seed set pre-expanded to ATP (see main text). If a metabolite is a reactant in all first reactions, the fold lineage is said to *emerge to use* (ETU) the metabolite, if this condition was met in 75% or more runs. If a metabolite is a reactant in some but not all of the first reactions, the fold lineage was classified as *first use* (FU) for the given metabolite, if this condition was met in 75% or more runs. If the 75% threshold is not met for either an emerge to use or first use classification, an emergence relationship was not assigned. In the case of molecular oxygen, no fold lineage falls into the unassigned category (**Figure 5C**). For fold lineages where none of the first reactions require the metabolite as a reactant across all runs, classification was based on the following criteria: L*earn to use* (LTU) if a fold lineage catalyzes a reaction using the metabolite as a reactant sometime later in the trajectory. *Indirect use* (IU) if a fold lineage never catalyzes any reaction that requires the metabolite, but does catalyze reactions that become unreachable when the metabolite is ablated from the network. *Never use* (NU) if a fold lineage never catalyzes reactions that either require the metabolite as a reactant, or become unreachable when the metabolite is ablated from the network. For any fold lineage, multiple metabolites can fall into the same category (*e.g.*, Rossmanns learn to use FAD and molecular oxygen), but each metabolite can only be related to a fold lineage by a single relationship (that is, fold lineage-metabolite pairs are uniquely classified). **Figure S19** presents a detailed classification of oxygen–fold lineage emergence relationships.

**Algorithm:** Enzyme-gated Network Expansion

```
Input: metabolic_reactions: set of allowed reactions, their reactants and products
       seed_compounds: set of starting metabolites
       reaction_to_rules: fold sets required to catalyze each reaction
       metabolic_folds: set of all folds associated with reactions in the metabolic model
Output: compound2iter, reaction2iter, rule2iter, fold2iter

Function expandOneStep(i, C, R, U, F):
    /* i : current iteration number                                    */
    /* C : dict of compounds and their iteration of discovery          */
    /* R : dict of reactions and their iteration of discovery          */
    /* U : dict of rules and their iteration of discovery              */
    /* F : dict of folds and their iteration of discovery              */
    i ← i + 1
    U_new ← U
    undiscovered_rules ← reaction_to_rules − {U.keys()}
    foreach rule ∈ undiscovered_rules do
        if all reactants of rule in C.keys() and
        all required folds of rule in F.keys() then
            U_new[rule] ← i ;                                       // new rule
            if reaction associated with rule ∉ R.keys() then
                R[reaction associated with rule] ← i ;              // new reaction
            C[c] ← i   ∀c ∈ {products of rule ∉ C.keys()} ;         // new compounds
    if U_new ≠ U then
        return i, C, R, U_new ;                                     // network expanded
    else
        return None ;                                               // no expansion

Function expandUntilStop(i, C, R, U, F):
    while expandOneStep(i, C, R, U, F) ≠ None do
        i, C, R, U ← expandOneStep(i, C, R, U, F)
    return i, C, R, U

Function getTopRankedfold(C, R, U, F):
    /* Rank folds                                                      */
    undiscovered_rules ← reaction_to_rules − {U.keys()}
    remainder_fold_sets ← sets of folds that would satisfy undiscovered_rules excluding {F.keys()}
    fold_utility ← dict()
    min_size ← 1
    while fold_utility = dict() do
        foreach s of size min_size in remainder_fold_sets do
            fold_utility[s] ← count(new rules added by s), if count > 0
        min_size ← min_size + 1 ;      // minimize the number of folds added simultaneously
    /* Select top-ranked fold(s)                                       */
    if count([s with max fold_utility[s]]) = 1 then
        top_ranked ← s
    else
        top_ranked ← choose one s randomly with max fold_utility[s] ;   // non-deterministic
    return top_ranked
```

```
Initialize:
iter ← 0
compound2iter ← dict()
compound2iter[c] ← iter    ∀c ∈ seed_compounds
reaction2iter ← dict()
rule2iter ← dict()
fold2iter ← dict()
/* Step 0:  Enzyme-independent expansion                                    */
iter, compound2iter, reaction2iter, rule2iter, fold2iter ← expandUntilStop(iter, compound2iter,
  reaction2iter, rule2iter, fold2iter)
while {F.keys()} ≠ metabolic_folds do
    /* Step 1:  fold Selection and Addition                                 */
    top_ranked ← getTopRankedfold(compound2iter, reaction2iter, rule2iter, fold2iter)
    fold2iter[f] ← iter + 1    ∀f ∈ top_ranked
    /* Step 2:  Standard Network Expansion                                  */
    iter, compound2iter, reaction2iter, rule2iter, fold2iter ← expandUntilStop(iter, compound2iter,
      reaction2iter, rule2iter, fold2iter) ;                    // defines metabolic stage
/* Output:  Complete enzyme-metabolism co-evolution trajectory              */
return compound2iter, reaction2iter, rule2iter, fold2iter
```

**Figure S1.** Pseudocode implementation of the enzyme-gated network expansion algorithm. In the code, each unique reaction–fold lineage set pair is referred to as a “rule” (**Table S7**). For simplicity, the main text frames the algorithm in terms of reactions — however, the fold lineage selection criteria consider rules as defined here, not reactions.

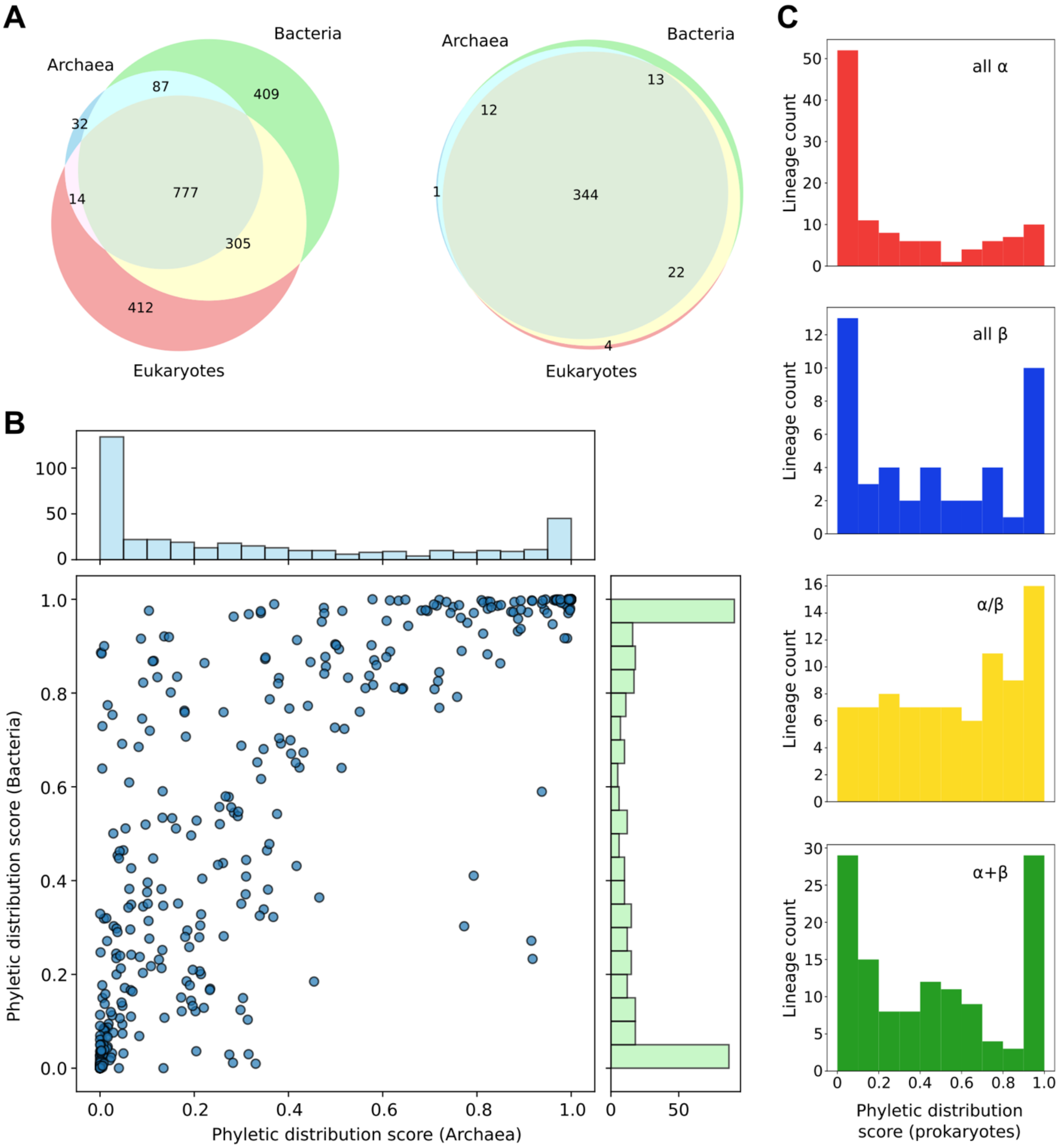


**Figure S2. A**. Distribution of all ECOD fold lineages (left, n = 2,230) and metabolic fold lineages (right, n = 396) in archaea, bacteria, and eukaryotes. Note that 228 fold lineages were not detected in any of the three domains of life. A fold lineage is considered present if it is detected in at least one genome. **B**. Phyletic distribution scores of metabolic fold lineages in archaea and bacteria. See **Methods** for a detailed explanation of the calculation. See **Table S1** for phyletic distribution scores of all ECOD fold lineages in archaea, bacteria, and eukarya (GTDB v.207, Eukprot v.3). **C**. Prokaryotic distribution score of fold lineages in all α, all β, α/β and α+β structural classes. The overall trend corresponds to previous studies (Abeln and Deane 2015) where α/β domains are more distributed.

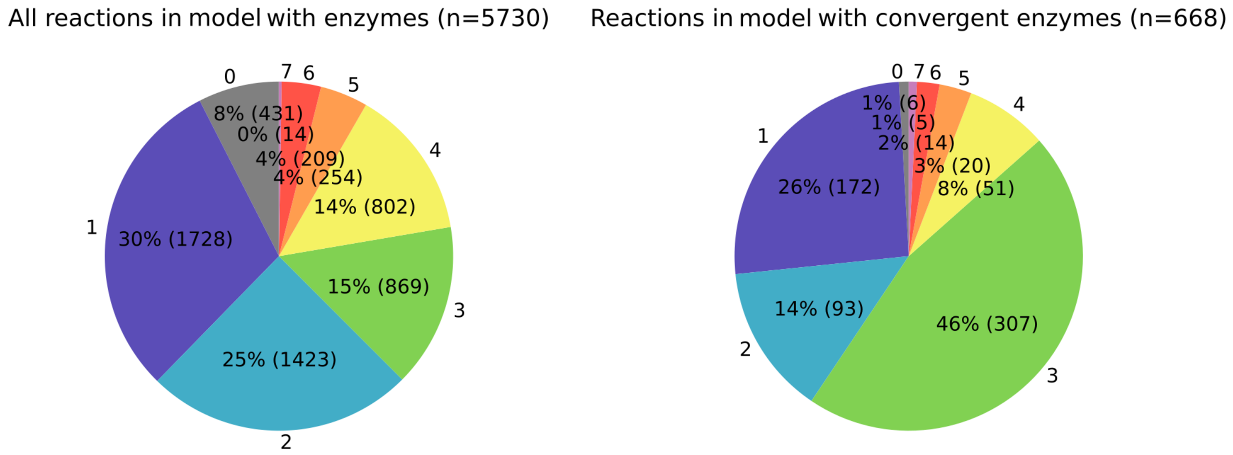


**Figure S3**. Reaction type breakdown of reactions in our model with enzymes (left) and the subset of reactions with convergent enzymes (right) where multiple mutually exclusive sets of fold lineage were mapped to the same reaction. The pie charts depict the proportion of reactions classified under each first-digit Enzyme Commission (EC) number. Approximately 12% (668/5,730) of enzyme-catalyzed reactions can be catalyzed by entirely distinct sets of fold lineages. It is worth noting that EC number 3 (hydrolase) is enriched in convergent enzymes.

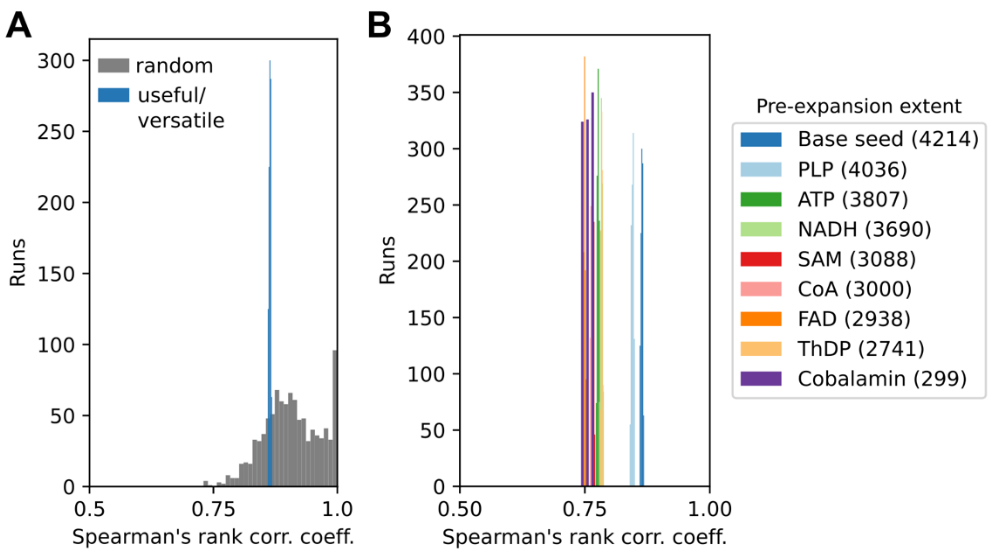


**Figure S4.** Compound discovery order is robust to fold lineage ordering. **A**. Spearman's rank correlation coefficient of standard versus enzyme-gated network expansion in which fold lineages were selected randomly (random) or based on the number of reactions that they satisfy at that point (useful/versatile). Note that the standard network expansion algorithm is deterministic. Although random fold lineage selection introduces significant variability compared to useful/versatile fold lineage selection, both approaches return compound discovery orders consistent with standard network expansion (useful/versatile: mean Spearman's rank correlation coefficient = 0.86; SD = 0.0019; random: mean Spearman's rank correlation coefficient = 0.91; SD = 0.057). Enzyme-gated network expansion refines, but does not significantly alter, the course of metabolic development inferred from standard network expansion — with the only exception of CoA, which is discovered early as a result of adding *L*-valine to the seed set. **B**. Compound discovery order of enzyme-gated network expansion with various degrees of seed set pre-expansion versus standard network expansion. 1,000 independent runs were calculated for each pre-expansion extent. Numbers in parentheses represent the number of compounds used for the comparison with standard network expansion, which excludes seed compounds from consideration.

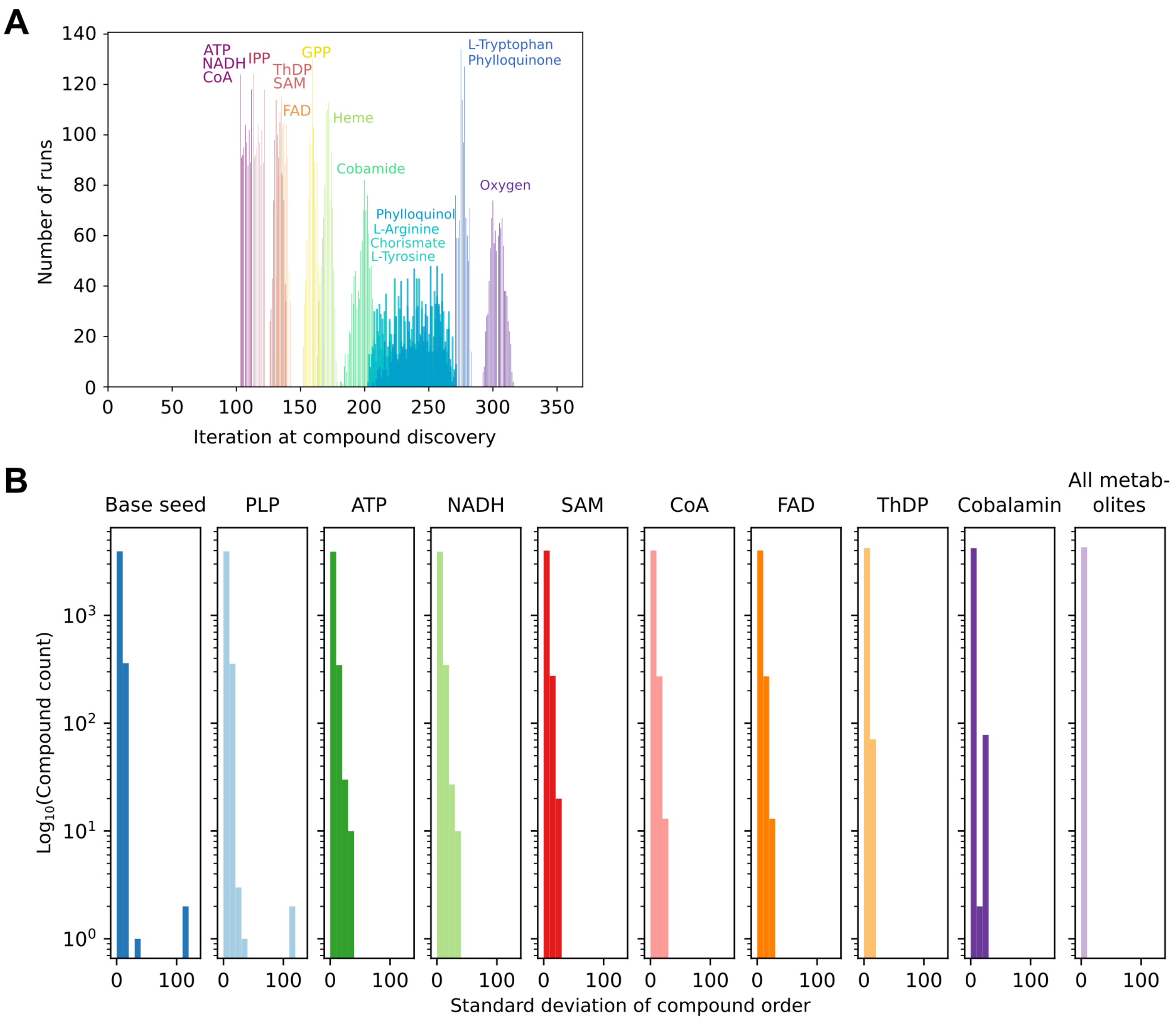


**Figure S5.** Compounds with variable discovery orders. **A**. Variability in the discovery order of key metabolites across 1,000 runs with no seed set pre-expansion. Variability is higher between iterations 200 and 270, which corresponds to the region where many equivalent single-reaction fold lineages (see **Figure S7**) are selected randomly. “Iterations” here and in subsequent supplementary figures include both the fold lineage addition steps and the intervening standard network expansion steps between fold lineage additions. **B**. Standard deviation of compound discovery order across 1,000 runs for various seed set pre-expansions. Standard deviation is zero for all compounds in the “All metabolites” pre-expansion because the simulation starts with all compounds in the seed set.

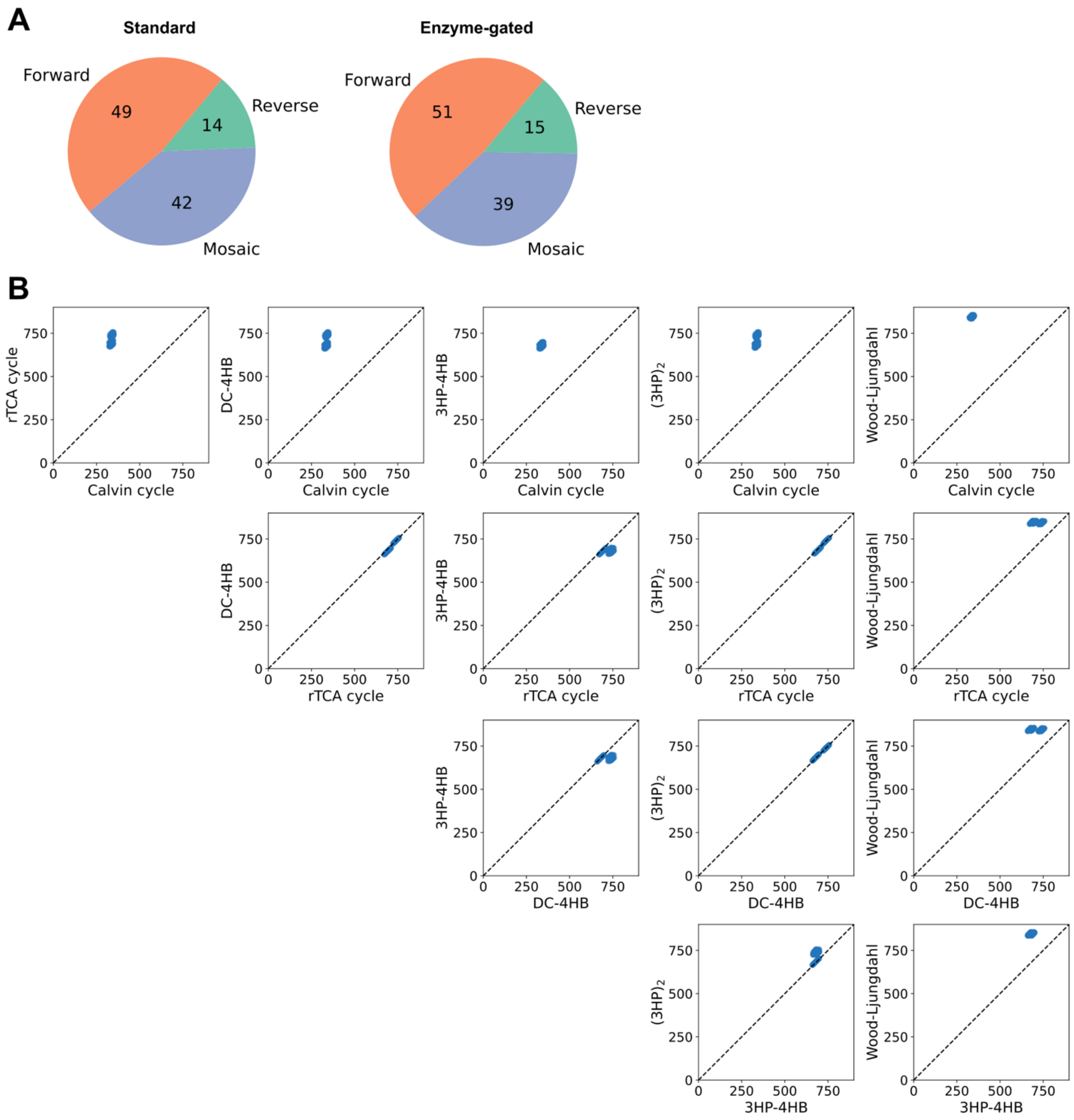


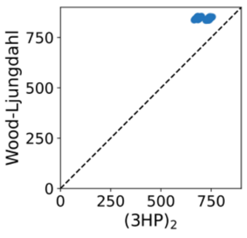


| Pathway name | Module ID | Mean | SD | Alternative name |
|---|---|---|---|---|
| Calvin cycle | M00165 | 337.6 | 6.7 | Reductive pentose phosphate cycle |
| 3HP-4HB | M00375 | 680.8 | 10.8 | Hydroxypropionate-hydroxybutylate cycle |
| DC-4HB | M00374 | 708.7 | 31.4 | Dicarboxylate-hydroxybutyrate cycle |
| $(3HP)_2$ | M00376 | 711.3 | 29.0 | 3-Hydroxypropionate bi-cycle |
| rTCA cycle | M00173 | 715.0 | 25.7 | Reductive citrate cycle |
| Wood-Ljungdahl | M00377 | 845.2 | 6.2 | Reductive acetyl-CoA pathway |

**Figure S6. A**. 105 linear metabolic pathways in the KEGG database were classified as emerging sequentially (forward or reverse) or non-sequentially (mosaic) based on the order of reaction discovery within each pathway (**Table S9**). The proportions between each mode of pathway emergence did not change significantly between standard network expansion (Goldford et al. 2024) and enzyme-gated network expansion using the base seed set. **B**. Emergence order of carbon fixation pathways. Scatterplots describe all-versus-all comparisons of the iteration of pathway completion for 6 carbon fixation pathways (Calvin cycle, 3HP-4HB, DC-4HB, $(3HP)_2$, rTCA cycle, Wood-Ljungdahl pathway). A pathway is considered complete when all of its constituent reactions have been discovered. Consistent with the standard network expansion results, enzyme-gated network expansion (base seed set, 1,000 runs) favors the early discovery of the Calvin cycle. The relative ordering of carbon fixation pathways likely reflects their differing cofactor requirements. Note that the base seed set used in these experiments assumes a geochemically supported rTCA-Wood-Ljungdahl pathway. The mean and standard deviation of pathway completion timing for each carbon fixation pathway are shown in the table.

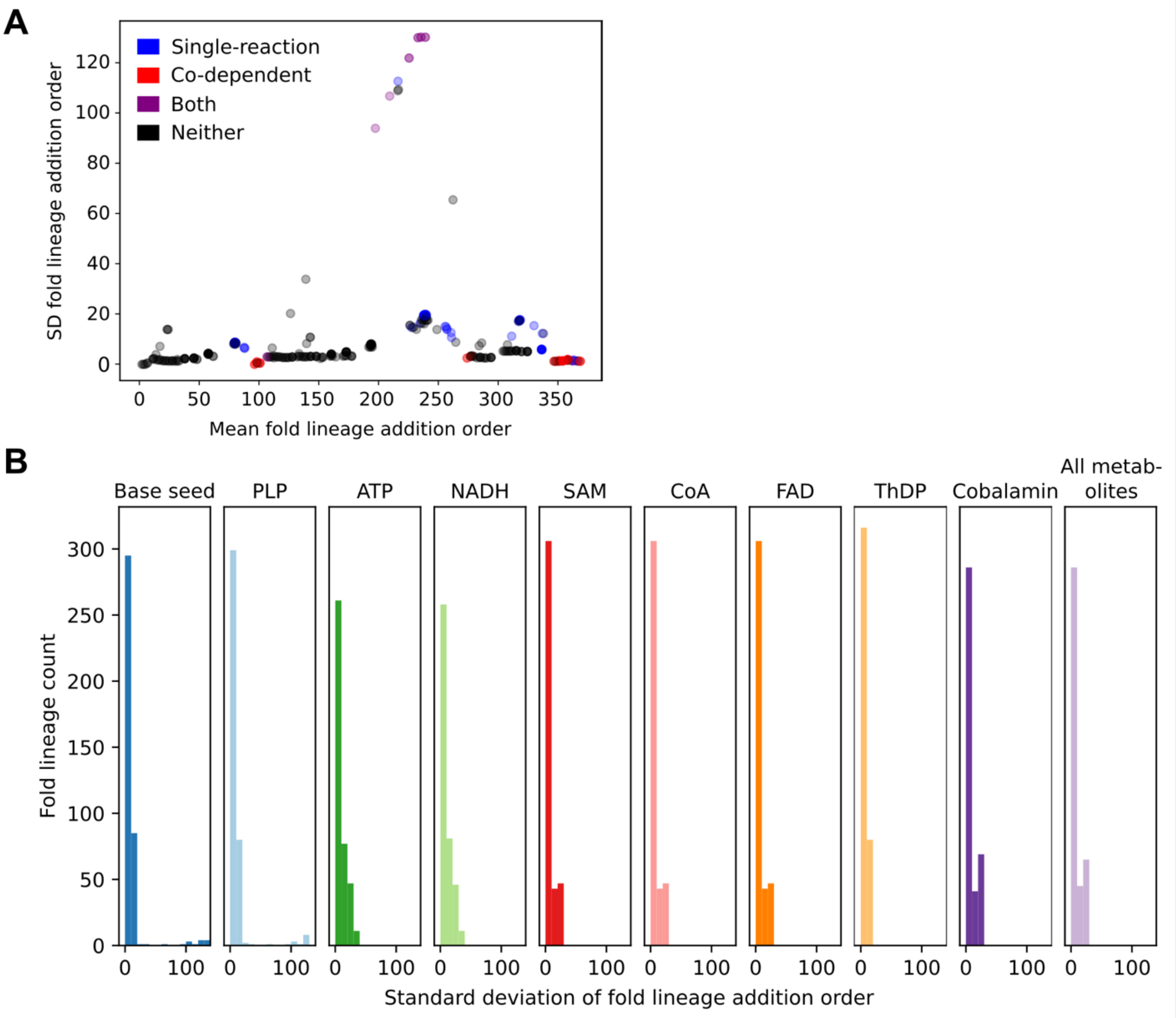


**Figure S7.** Standard deviation of fold lineage addition order is low for the vast majority of fold lineages in all levels of pre-expansion. **A**. Standard deviation versus mean fold lineage addition order for 1,000 runs with no seed set pre-expansion (base seed set). Standard deviation is higher for 2 types of fold lineages: "single-reaction" fold lineages (blue) that introduce only one new reaction upon addition in 50% or more runs, and highly "co-dependent" fold lineages (red) that require multiple fold lineages to be added simultaneously for further expansion in ≥50% of runs. **B**. Histograms of the standard deviation of fold lineage addition order for all fold lineages in simulations with various seed set pre-expansion extents, 1,000 runs each. Standard deviation is low for the vast majority of fold lineages, with the exception of single-reaction and co-dependent fold lineages, particularly in runs with little or no pre-expansion.

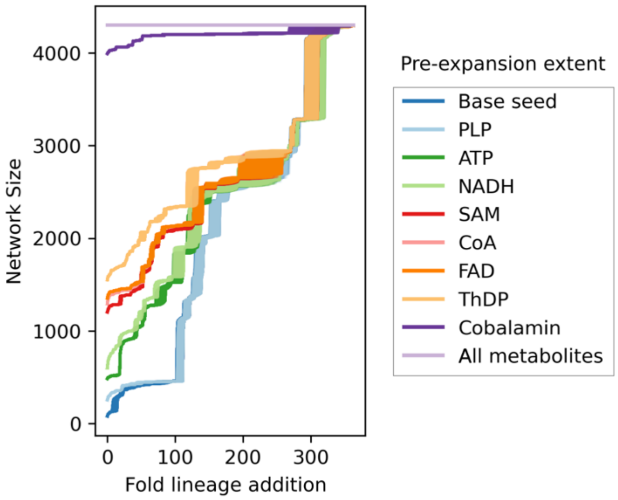


**Figure S8.** Network expansion trajectories for varying degrees of seed set pre-expansion (1,000 runs each).

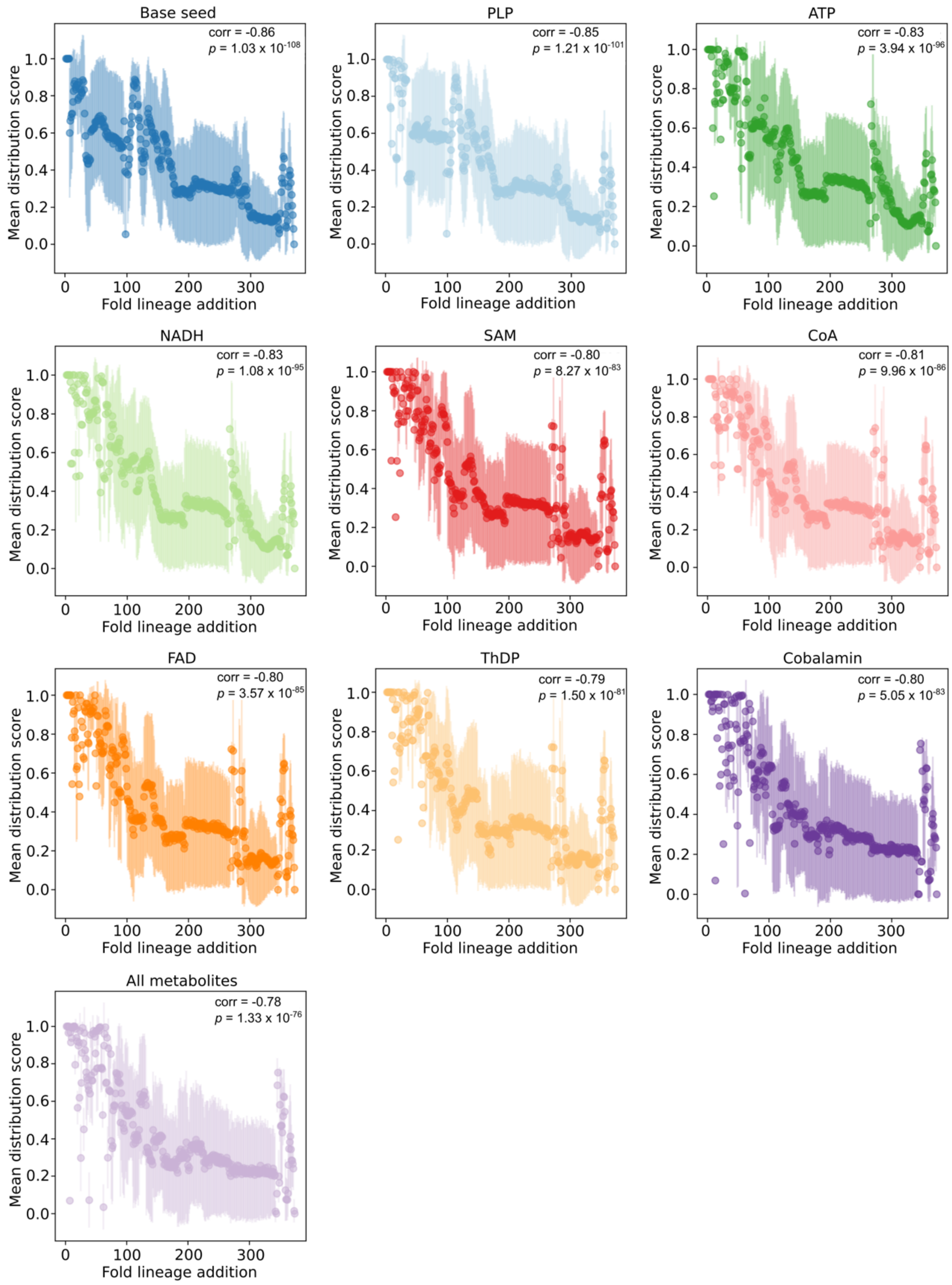


**Figure S9.** Average prokaryotic distribution at each fold lineage addition step across 1,000 runs for varying degrees of seed set pre-expansion. The correlation between fold lineage addition order and prokaryotic distribution scores (see **Methods**) persists regardless of the extent of pre-expansion. P-values from Spearman's rank correlation.

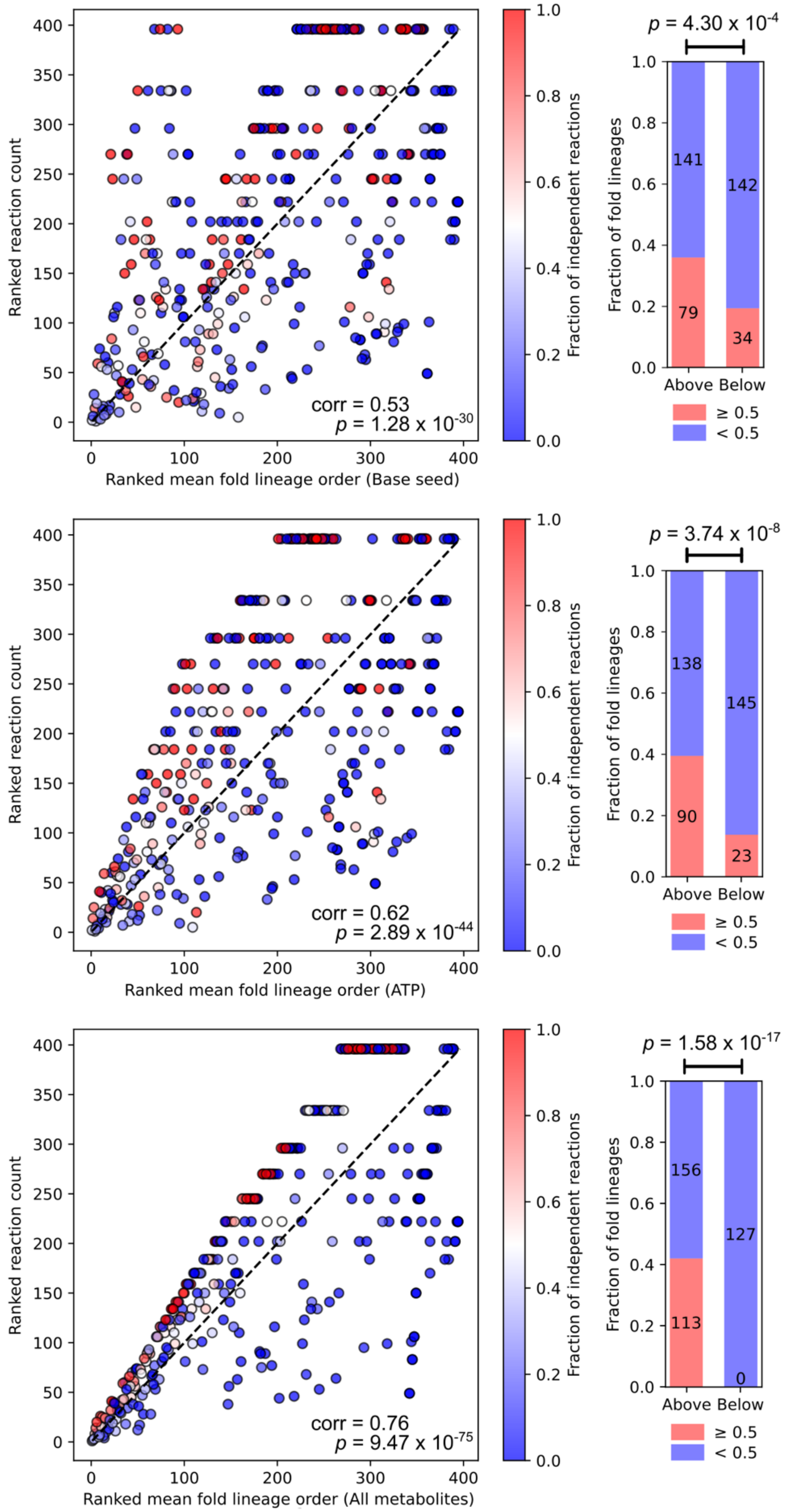

Ranked reaction count
Ranked mean fold lineage order (Base seed)
corr = 0.53
p = 1.28 x 10-30
Fraction of independent reactions
p = 4.30 x 10-4
Fraction of fold lineages
141
142
79
34
Above Below
≥ 0.5
< 0.5
Ranked reaction count
Ranked mean fold lineage order (ATP)
corr = 0.62
p = 2.89 x 10-44
Fraction of independent reactions
p = 3.74 x 10-8
Fraction of fold lineages
138
145
90
23
Above Below
≥ 0.5
< 0.5
Ranked reaction count
Ranked mean fold lineage order (All metabolites)
corr = 0.76
p = 9.47 x 10-75
Fraction of independent reactions
p = 1.58 x 10-17
Fraction of fold lineages
156
127
113
0
Above Below
≥ 0.5
< 0.5

**Figure S10.** Greater extents of seed set pre-expansion prioritize fold lineages with high reaction counts that can catalyze reactions independently. For three degrees of pre-expansion (base seed, ATP, All metabolites), the ranked mean addition order is plotted against the ranked reaction count for each fold lineage. Datapoints are colored according to the fraction of reactions associated with a fold lineage that it can catalyze independently, without the help of a co-domain. With no pre-expansion (top), some fold lineages are added early despite their low reaction count while others are added late despite their high reaction count. This results from the underlying metabolic network and its dependency structure prevents fold lineages from being added purely based on their reaction count, instead selecting the useful/versatile fold lineage based on the available compounds it can use and its ability to cooperate with pre-existing fold lineages. Starting from a fully expanded seed set where all compounds have already been discovered (bottom) strengthens the rank correlation between reaction count and addition order (Spearman's rank correlation coefficient = 0.73). This is especially true for fold lineages that can independently catalyze reactions. Fold lineages capable of independently catalyzing ≥ 0.5 of their associated reactions increasingly concentrate above the diagonal, indicating that their discovery order rank is lower than their reaction count rank (bar plots, right). P-values for differences in fold lineage fractions calculated using a chi-square test.

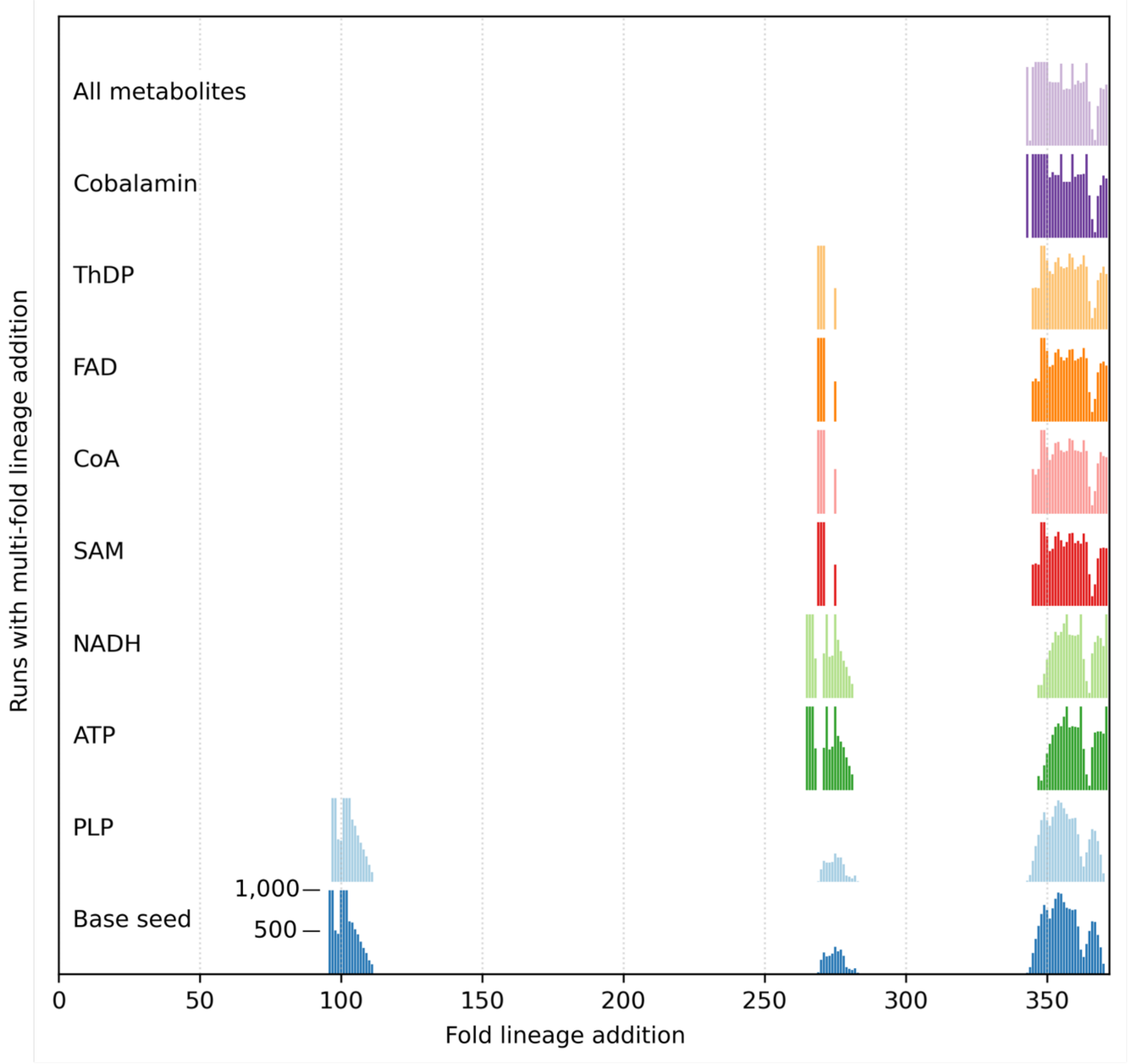


**Figure S11.** Periods of multi-fold lineage discovery for various seed set pre-expansion extents. Both the base seed (80 compounds) and PLP (255 compounds) seed sets require the early emergence of multiple domains simultaneously around 100 fold lineage additions to produce ATP. The existence of obligatory multi-fold lineage addition postpones the production of ATP until 100 fold lineage additions or more, even though many of the already-discovered fold lineages are known to extensively use ATP, such as the P-Loop NTPases and ATP grasps. These data demonstrate that the timing of multi-fold lineage addition is consistent across 1,000 simulations for each pre-expansion extent.

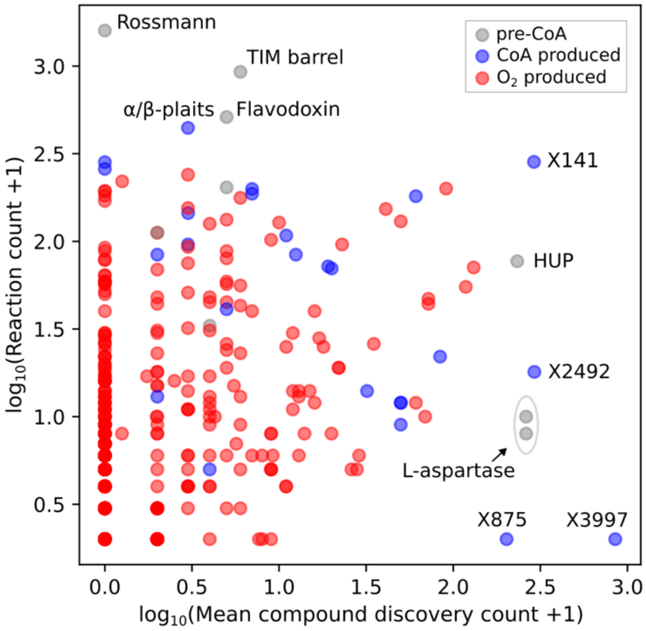


**Figure S12.** Burst analysis. Large bursts in compound discovery (x-axis) are often triggered by the addition of fold lineages that uniquely catalyze the synthesis of highly connected hub metabolites or their precursors. These fold lineages are not necessarily associated with many reactions (y-axis) — in fact, X875 and X3997, which unlock chorismate and chlorophyll (and subsequently oxygen), only have a single associated reaction. Burst-inducing fold lineages and their biosynthetic associations are as follows: X141 (Geranyl diphosphate), HUP (NAD), X2492 (Flavin precursor), *L*-aspartase (purine precursor), X875 (chorismate), X3997 (chlorophyll).

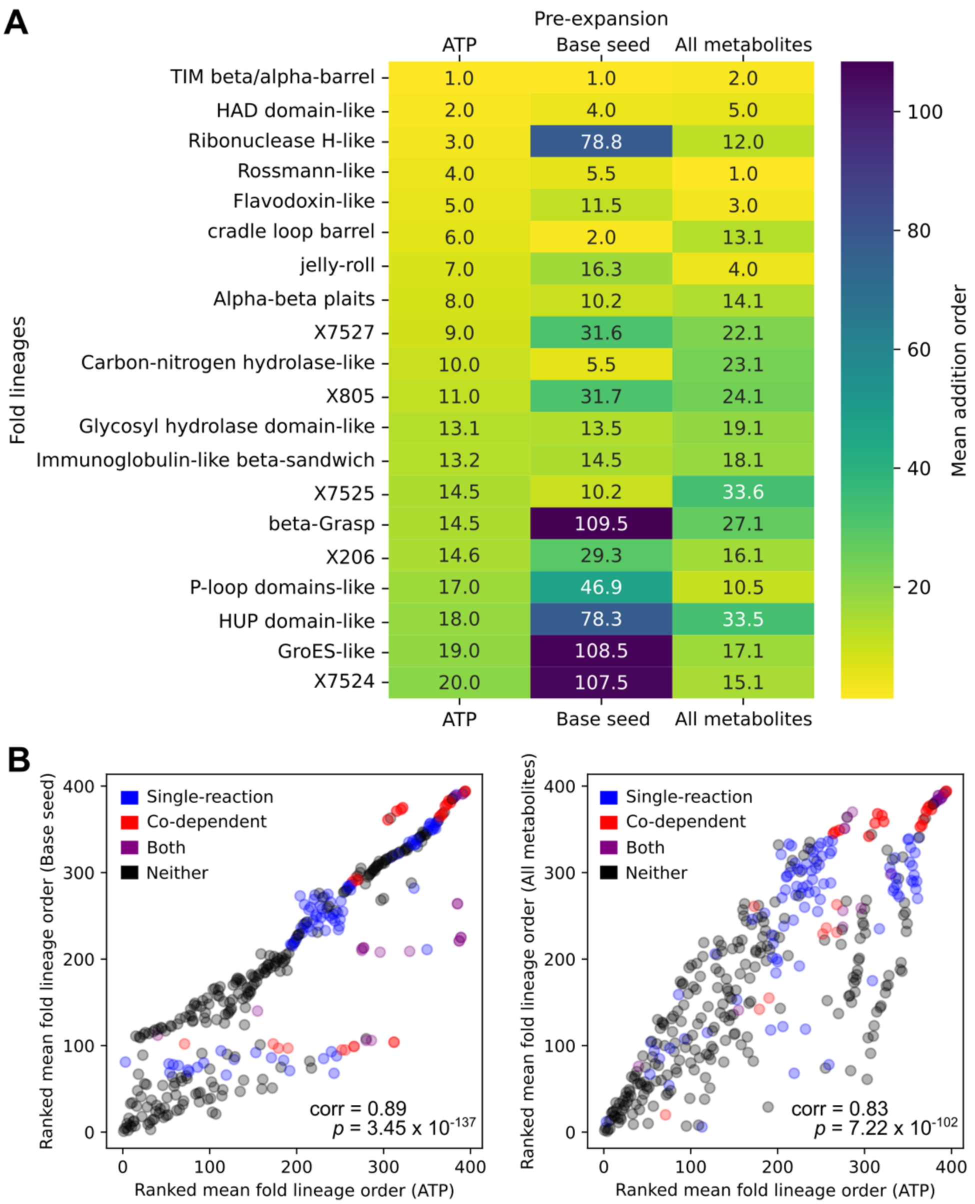


**Figure S13. A**. Comparison of mean fold lineage addition orders of the first 20 fold lineages with seed set pre-expansion to ATP with those from no pre-expansion (base seed) and full pre-expansion (All metabolites). Although differences between pre-expansions exist, fold lineage orders in each case are highly correlated for most early fold lineages. **B**. Comparison of ranked mean fold lineage orders of all metabolic fold lineages from ATP, base seed, and all metabolites pre-expansions. Large deviations in fold lineage orders between various pre-expansions are often attributed to “single-reaction” fold lineages (blue) that introduce only one new reaction upon addition in 50% or more runs, and “co-dependent” fold lineages (red) that require multiple fold lineages to be added simultaneously for further expansion in 50% or more runs. P-values from Spearman’s rank correlation tests. See also **Figure S7**.

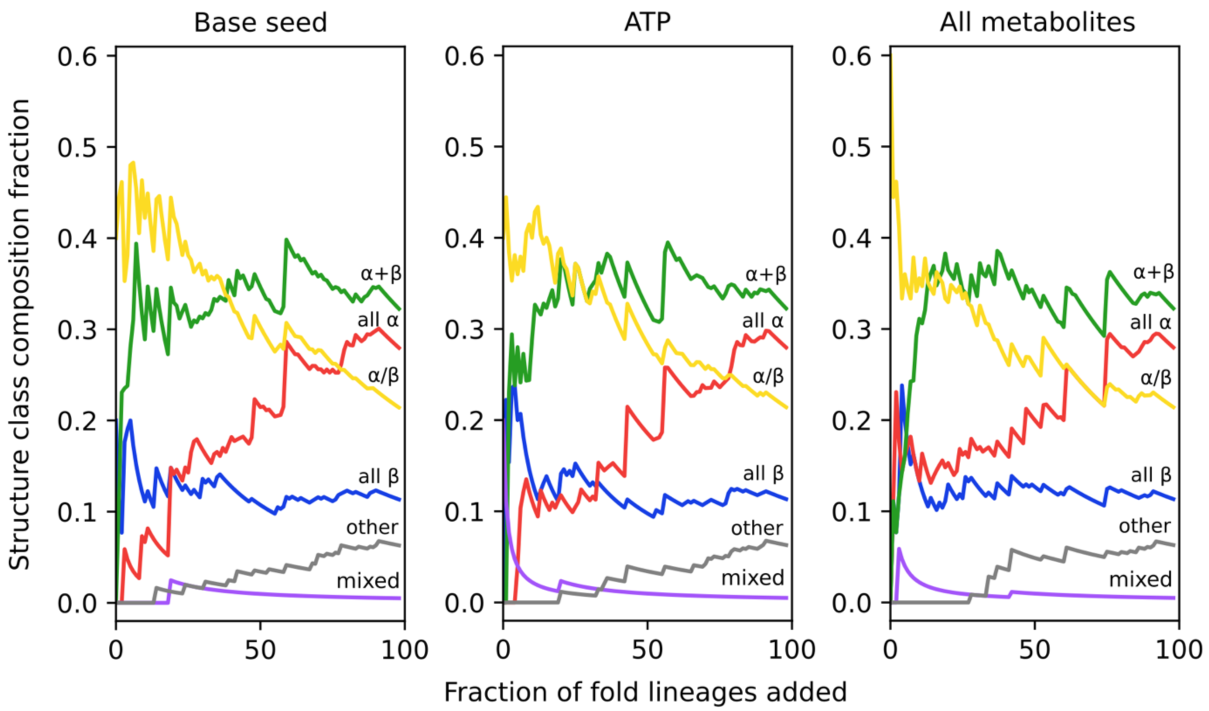


**Figure S14.** Protein structure class discovery. Averaged over 1,000 runs. Comparison between simulations with different seed set pre-expansion extents suggest the early dominance of α/β class is a robust feature regardless of when proteins are introduced as the dominant catalysts in metabolic evolution.

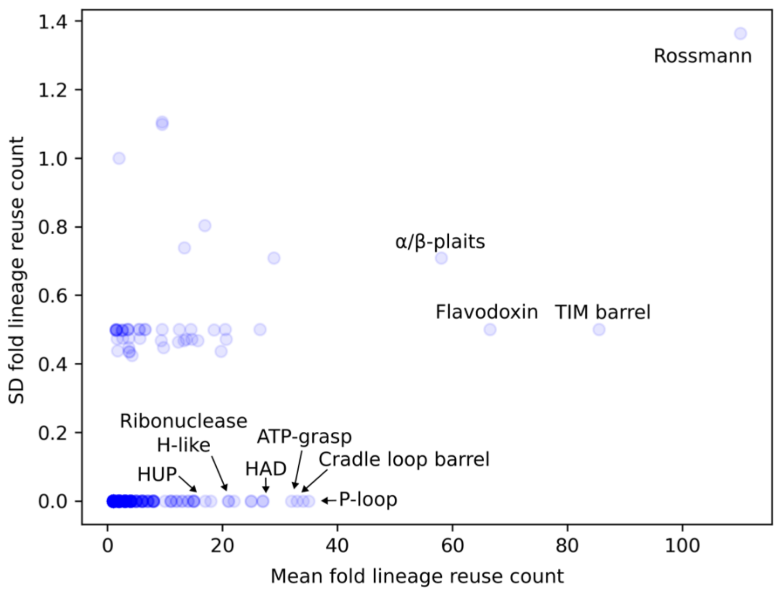


**Figure S15.** Mean versus standard deviation of the number of iterations where each fold lineage is reused, averaged across 1,000 runs of the simulation with pre-expansion to ATP. A fold lineage is said to be reused when it catalyzes a new reaction or reactions at an iteration later than its emergence. 54% of metabolic fold lineages (216/396) are recruited at multiple points along the trajectory. Standard deviation of reuse count is low for all fold lineages. Rossmann (X-group 2003), TIM barrel (X-group 2002), and flavodoxin (X-group 2007) are the most reused fold lineages.

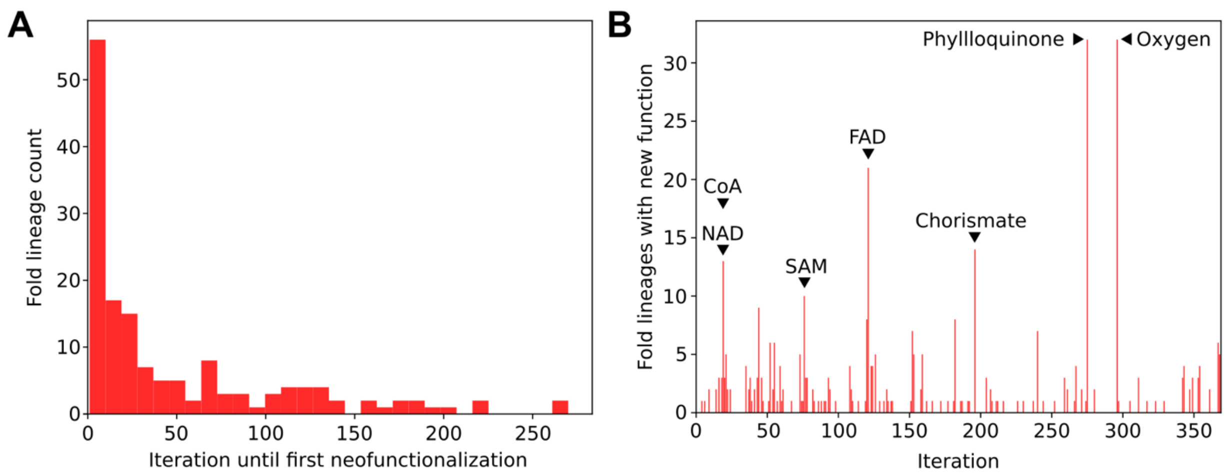


**Figure S16. A**. Number of fold lineage additions until first neofunctionalization, defined here as discovery of a reaction with a novel 2-digit EC number. The majority of fold lineages (90/156 or 58%) perform a new function within 30 fold lineage additions after emergence. **B**. Number of fold lineages that undergo neofunctionalization at each iteration. Both panels A. and B. are based on the representative run shown in main text **Figure 4**.

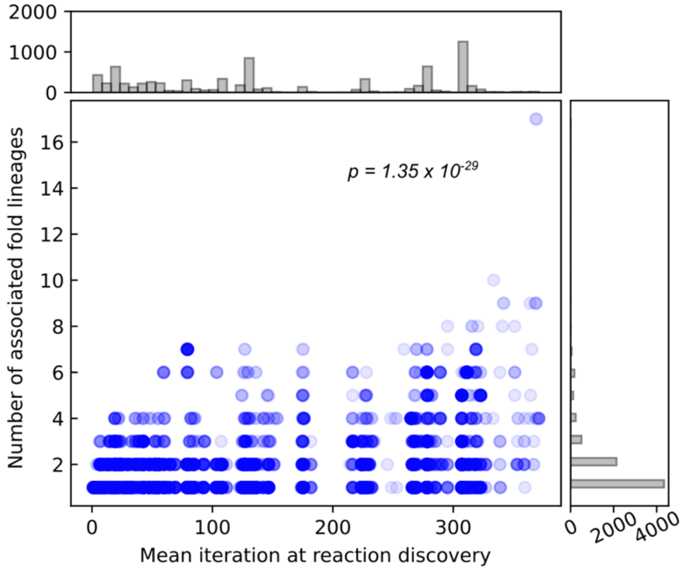


**Figure S17.** Fold lineage set complexity generally increases across network expansion. For each reaction, its mean iteration of discovery and the number of associated fold lineages is plotted. P-value calculated by the Pearson correlation test.

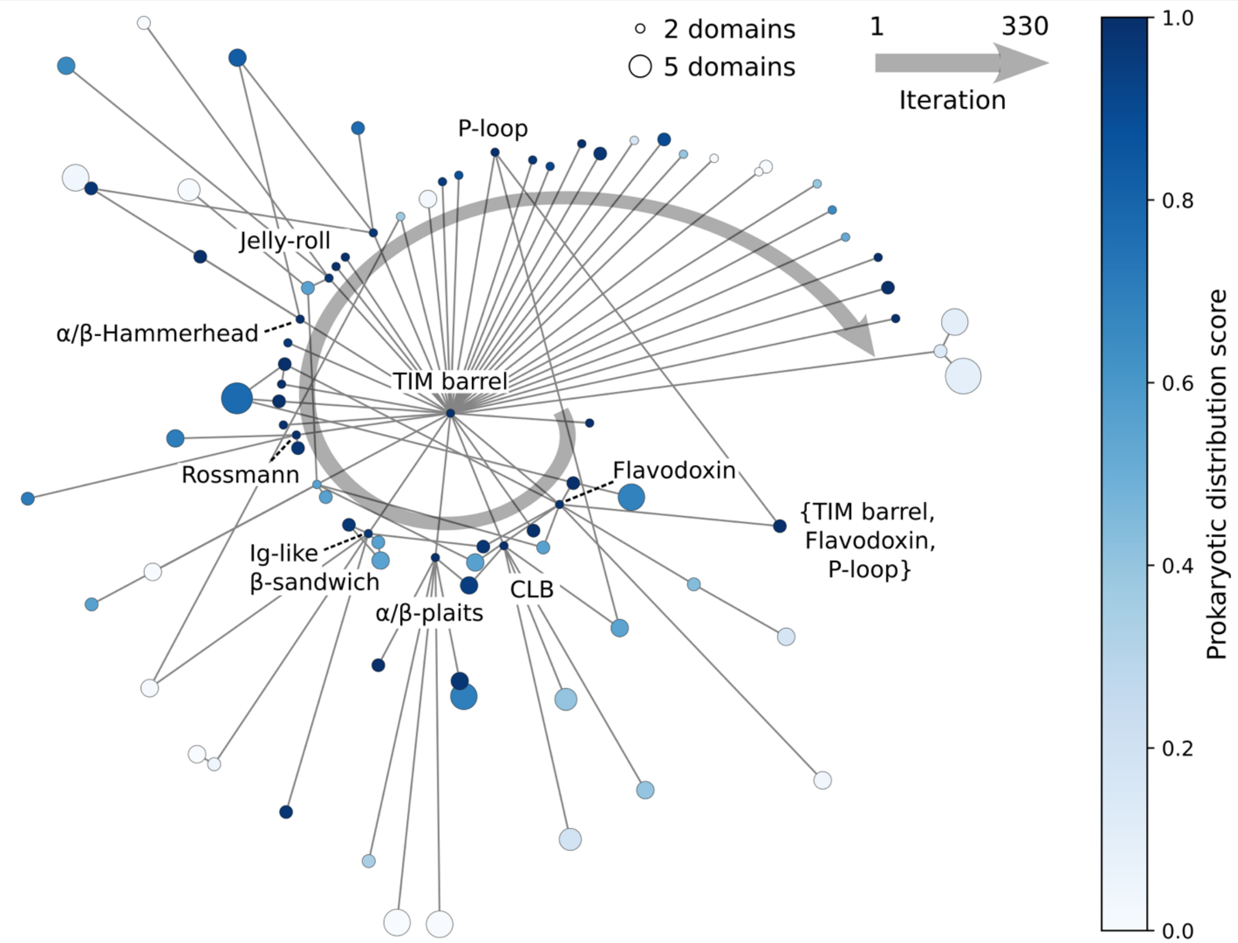


**Figure S18.** Domain accretion history of TIM barrel (X-group 2002) along the representative run shown in **Figure 4**. The TIM barrel, which is the first fold lineage to emerge, is placed at the center of the figure. Each node represents a novel domain accretion event in which the TIM barrel jointly catalyzes a reaction with one or more other fold lineages. The distance of each node from the center TIM barrel node corresponds linearly to the first iteration where that fold lineage set catalyzed a reaction. Note that other fold lineages or fold lineage sets may have catalyzed these reactions first. Nodes in the outer layer are connected to those in the inner layer if they form a super set as in the case of {X2002, X304, X230} and {X2002, X304}. Node sizes correspond to the number of fold lineages in the fold lineage set. The spiral indicates the relative order of the first layer of domain accretion around the TIM barrel spanning iterations 1 to 330. The positions of the outer-layer nodes were adjusted to improve readability while precisely retaining the distance from the center. The colormap reflects the prokaryotic distribution score of each fold lineage set, calculated based on the co-occurrence of all fold lineages within the set, rather than individual fold lineages.

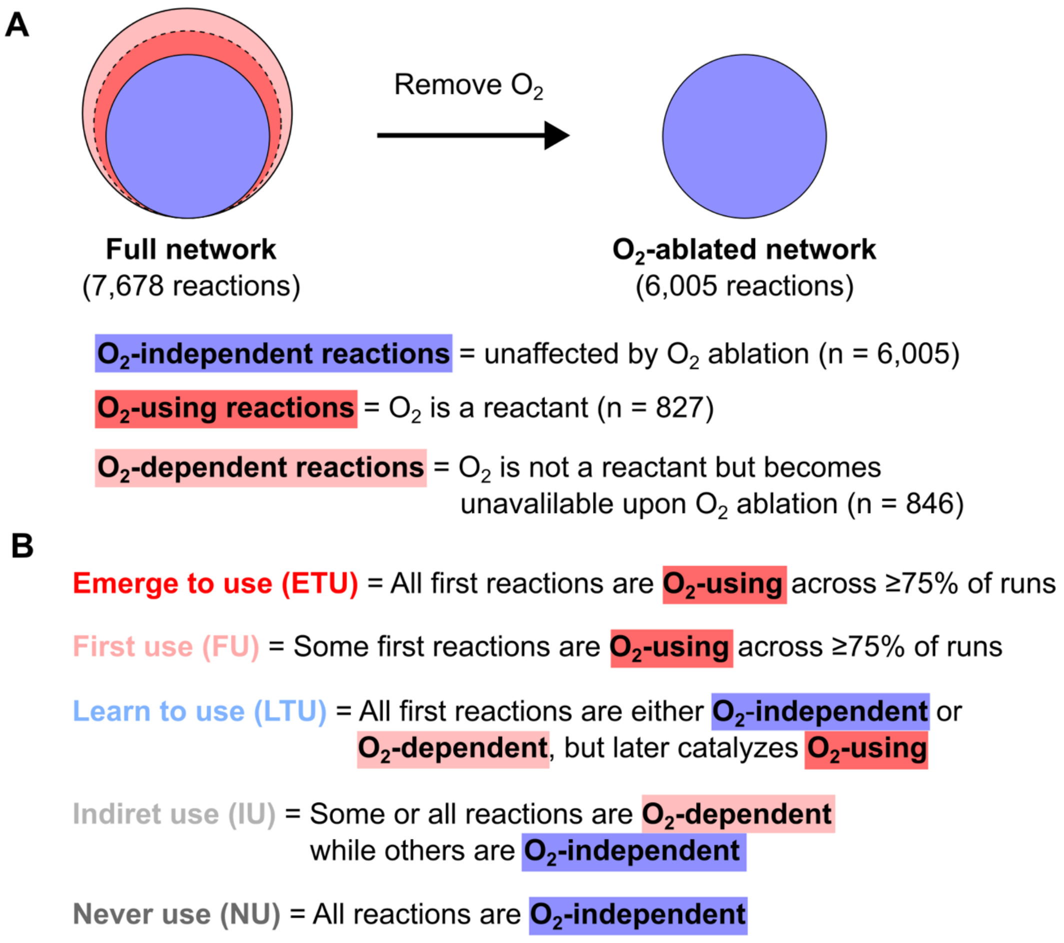


**Figure S19.** Fold lineage-oxygen emergence relationships. **A**. Ablating $O_2$ from the network and performing expansion prevents the discovery of 1,673 reactions, of which 827 directly use $O_2$ ($O_2$ is a reactant; *$O_2$-using*) and 846 depend on $O_2$ but do not directly use $O_2$ (either $O_2$ is a product, or one or more reactants are reachable only when $O_2$ is present; *$O_2$-dependent*). The remaining 6,005 reactions are reachable without $O_2$ (*$O_2$-independent*). **B**. For emerge to use (ETU) fold lineages, all reactions catalyzed by that fold lineage in the first network expansion step after fold lineage addition are $O_2$-using reactions, and this property is consistent across ≥75% of runs. For the first use (FU) fold lineages, $O_2$-using reactions are among the first reactions catalyzed by this fold lineage while reactions that do not involve $O_2$ (either $O_2$-independent or $O_2$-dependent) are also present. As before, consistency across ≥75% of runs is required. For FU fold lineages, we cannot resolve whether the $O_2$-using reaction came first, whereas with ETU this ambiguity does not exist. Learn to use (LTU) fold lineages do not catalyze *$O_2$-using* reactions when added, but later learn to catalyze one or more *$O_2$-using* reactions. Indirect use (IU) fold lineages must catalyze at least one *$O_2$-dependent* reaction, but not catalyze any $O_2$-using reactions. Never use (NU) fold lineages only catalyze *$O_2$-independent* reactions.

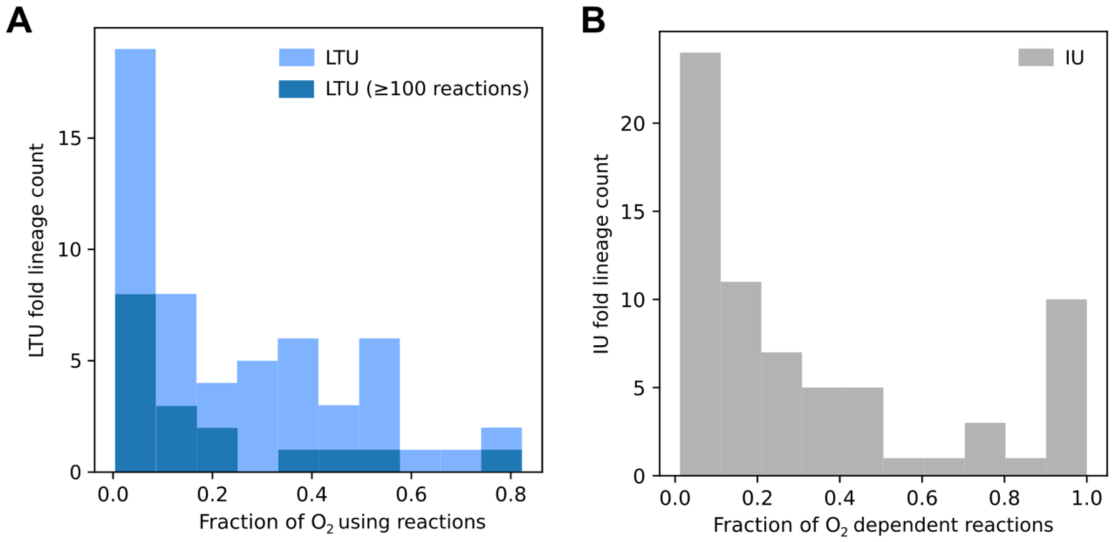


**Figure S20. A**. For most learn to use (LTU) fold lineages, $O_2$-using reactions constitute a small fraction of their total number of associated reactions. 65% (17/26) of versatile fold lineages (those with ≥100 associated reactions) are LTU. **B**. Likewise, $O_2$-dependent reactions constitute a small fraction of their total number of associated reactions for indirect use (IU) fold lineages.
